# Clustered heterogeneity, spiking history and efficient silencing maximize mutual information in adaptive time encoding

**DOI:** 10.64898/2026.09.09.750511

**Authors:** Raphaël Lafond-Mercier, Leonard Maler, André Longtin

## Abstract

Neurons involved in a computation are remarkably diverse. They display a range of thresholds, time scales and spiking history dependence, and often exhibit adaptation that enhances computational power by concentrating spikes at transient stimuli. We seek basic organizational principles for coding with cellular heterogeneity by focusing on the encoding and retrieval of inter-event time interval sequences by adaptive neurons. We formulate the general input-output mutual information maximization problem for parallel neurons with a fading memory of past intervals. The solution reveals an unexpected multi-variate heterogeneity that is tailored to encode information efficiently. Gradient ascent on mutual information produces a number of parameterized clusters bounded by the sequence length. Correlations between parameters emerge naturally. The predicted covariation of adaptation time with spiking history strength, and the optimal combination of history and non-history cells, agree with experiments. Threshold optimization yields a continuum of time constants and thresholds within each cluster. Strikingly, division of labour between neurons arises where subsets respond selectively to different interval ranges. This creates cell-specific interval ranges that tile reasonably well a uniform prior of intervals, and almost perfectly a scale-free prior, producing a code that is efficient in cell number and reduces metabolic cost by limiting spike rates.

## 1 Introduction

Biophysical heterogeneity of neurons within neural networks is omnipresent. Even when cells seemingly perform the same computation, their intrinsic dynamics, time constants and excitability can differ [1]. Heterogeneity has in fact been shown to support healthy brain activity [2]. However, fundamental organizational principles underlying the distribution of cell parameters are still needed in spite of much recent progress [3–5]. Metabolic constraints [6–8] argue for limiting firing rates, e.g. by drawing on a distribution of thresholds and firing fatigue, i.e. adaptation. How is threshold heterogeneity coordinated with variations in other parameters across a neural population to optimally convey information about stimuli?

We frame this general theoretical problem in the context of encoding sequences of intervals between discrete events on the scale of seconds to minutes given the vast interest in this topic [9–15]. Models of biologically plausible single cells [16–18] and networks [19, 20] have been put forth for encoding intervals in the hippocampus, and in the medial (MEC) and lateral (LEC) entorhinal cortex where “time cells” have been found [21–24]. These cells are proximal to spatially tuned cells named “place” and “grid cells” [25, 26]. Time cells tile time much like place and grid cells tile space, activating only after a specific amount of time has elapsed following an event. Both spatial and temporal information are important for path learning and episodic memory [27–29]. LEC even marks “events” through abrupt accelerations visible in low-dimensional trajectory projections in firing rate space [30]. Machine learning models such as recurrent neural nets (RNNs) [31–35] and reservoir computers [10, 18] have been trained to exhibit time cell properties. Their power lies in their connectivity, yet do not account for circuitry constraints nor spike rate adaptation seen in many hippocampal cell types [36, 37]. Adaptation has been suggested to perform predictive coding [38] and to enhance computational power of artificial and neuromorphic nets [39, 40].

One prominent modelling direction sees time cells implementing an approximate inverse Laplace transform associated with leaky integration and adaptation [41]. An ensemble of such neurons with different time constants could then drive another population that synthesizes the delayed responses seen in time cells, leading to scale invariance with a logarithmic range of time constants as seen in rat time cells [42]. Experimentally observed scale-invariant behaviour [43] could therefore be supported by such neurons in a biologically plausible manner, e.g. using simple gain control [44]. Neurons closely resembling these proposed “Laplace” cells were found in a thalamus-like region of weakly electric fish [45]. They signal whole body encounter events with environmental objects. Their time coding uses the number of spikes fired during an encounter, which increases with the time since the last encounter. Recovery-from-fatigue sees firing resources that were partially depleted at an event be replenished with a cell-specific “adaptation” time constant on the order of 0.5 to many seconds, much like a latent leaky integrator. Modelling shows that recovery can encode intervals using latent adaptation dynamics that control noisy (Poisson) firing [45]. These spikes then target the pallium - the highest brain level of vertebrates.

Theory shows that such adaptive encoding of sequences requires parametric heterogeneity [46]; establishing what heterogeneity *enables*; but importantly, the *optimized parametrization is unknown*. Moreover, three observed neural properties were omitted: a heterogeneous response threshold (RT) which silences neurons until enough resources have recovered, an external working memory (WM) which retains some information about past responses, and spiking history. This latter mechanism goes beyond the Laplace framework by accounting for the range of adaptation resetting after a spike [45, 46], and thus for how spiking (i.e. encounter) history could help decode sequences. Networks with such adaptation have been analyzed in other contexts using quasi-static approximations [47]. Here “adaptation memory” is a key organizer of heterogeneity.

We derive optimal distributions of recovery time, RT and history-dependence using mutual information (MI) between adaptive responses and intervals. Our previous work with Fisher information used specific interval sequence examples to retrieve sequence information from only the last population response. Here, through theory and intensive computational work, the full MI of sequences is optimized for different WM lifetimes. Gradient ascent converges to a parameter distribution that produces maximally informative responses given all possible (short) sequences consistent with the interval prior. This leads to sub-populations or “clusters”. Including RT in the optimization produces clusters tuned to certain intervals that together tile the prior. Without WM, the number of clusters equals the number of intervals to encode. Adaptation times depend on the prior, occur in clusters, with WM reducing cluster number. Our results derive from first principles efficient heterogeneity rules in adaptive neurons, some of which actually manifest in experiments [45].

## 2 Results

In this section, we proceed with defining the adaptation mechanism and how it generates responses. We introduce a model of working memory where noisy recalls of past responses is available for inference. We then show how sequences responsible for a set of observed responses and recalls can be recovered with the help of maximum likelihood estimation, noting the shortcomings of this method and therefore justifying the use of mutual information for a better analysis of the model. We then show how parametric heterogeneity emerges from the MI maximization and decompose the information content to reveal the mechanism behind the efficient encoding of time intervals in a sequence of consecutive events.

### Model definition

We start by defining the adaptation and threshold mechanism used to encode time sequences. For the entirety of the article, we use subscripts to label events and superscripts to label neurons or populations of identical neurons on non-boldface variables (i.e. scalar values). We model single-neuron resource availability with the variable *x* which is partially depleted during an encounter with an object [45, 48], and otherwise exponentially relaxes back to *x* = 1. This type of encounter is considered an event whose duration is much shorter than the time elapsed between encounters. During event *n*, the amount of available resources for the firing of cell *j* is given by

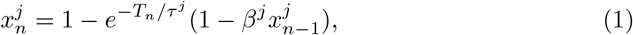

where *T*_*n*_ is the time elapsed between event *n* − 1 and event *n, τ* ^*j*^ is the recovery rate of the resources or “adaptation time constant”, and *β*^*j*^ ∈ [0, 1] is the remaining fraction of available resources after an event, which describes history dependence of the response of a cell, the “adaptation memory”. An example of the evolution of *x* is shown in Figure 1a.

**Fig. 1:**
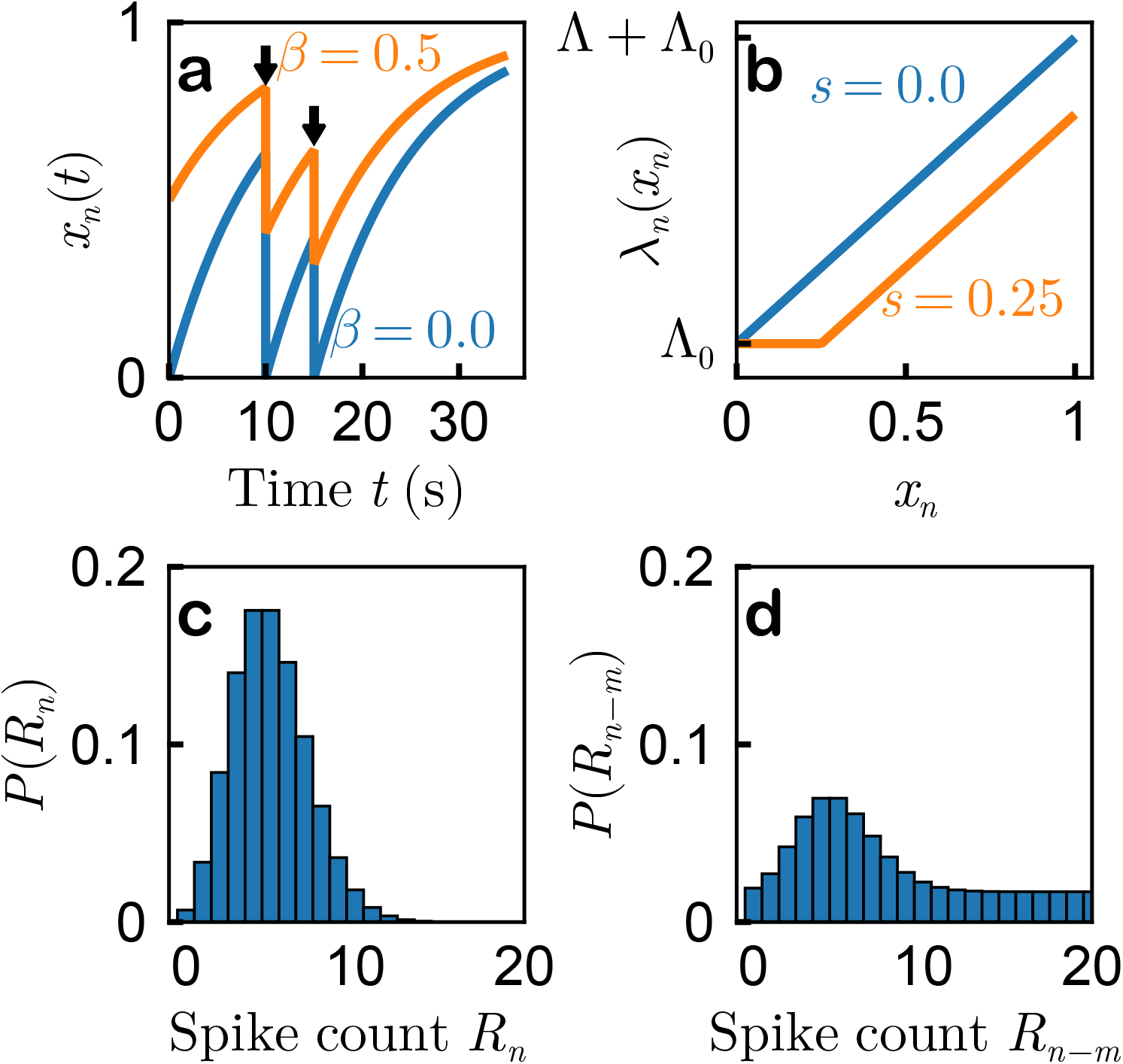
Modelling response adaptation and threshold with working memory. **(a)** Available resources *x* for an adaptive cell recover exponentially with time constant *τ*. After an event (black arrow), a fraction *β* of resources remains (see Equation (1)), as seen experimentally [45]. **(b)** A linear rectification is applied to the available resources *x* to retrieve a firing parameter *λ* during an event (see Equation (2)). A response threshold (RT) *s* keeps the firing parameter at baseline rate Λ_0_ when *x < s*. **(c)** The firing parameter *λ* is used to generate *R*_*n*_ spikes during event *n* with a Poisson distribution (see Equation (3)). **(d)** Working memory allows for the recall of responses that resulted from past encounters. When recalling previous response *R*_*n*−*m*_ during event *n*, the original Poisson distribution is combined with a uniform distribution to simulate the fading of past responses (see Equation (5)). This models downstream circuitry that receives adaptive responses which are corrupted by noise, stored in WM and recalled in the future.

During an event *n*, a firing parameter is generated for cell *j* and is given by

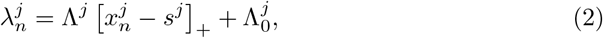

where Λ^*j*^ is the gain, 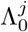 is the baseline firing parameter and *s*^*j*^ ∈ [0, 1] is the response threshold (RT) parameter. [·]_+_ is a rectified linear transformation: only cells with a resource variable 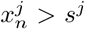 will have a rate that exceeds the baseline 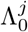. Examples of this rectification are shown in Figure 1b with (orange) and without (blue) RT.

The firing parameter is then used to generate the response of a neuron *j* during event *n* with a Poisson distribution:

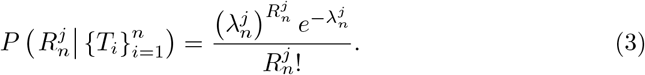

Real time cells are usually less variable than Poisson, which implies that our coding analysis pertains to a worst case scenario. An example response distribution is shown in Figure 1c. For a full network of *P* neurons, the distribution of responses is conditionally independent and is given by:

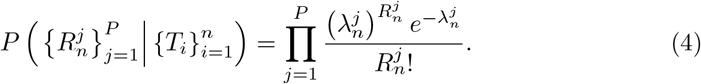

We omitted to write 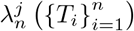 explicitly in Equations (3) and (4) to lighten the notation, but this quantity does depend on the sequence of time intervals.

Finally, working memory (WM) is modeled through the recall of past responses. During event *n*, the distribution of the recalled past response 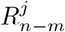 is given by a mixture of the original Poisson distribution and a uniform distribution:

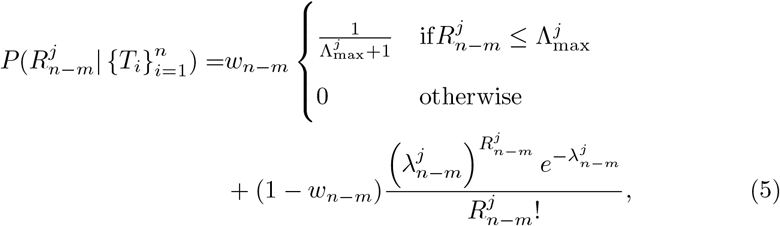

where *w*_*n*−*m*_ = min 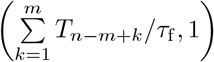 is the weight given to the uniform distribution which grows linearly with the time elapsed since event *n* − *m* until a maximum *τ*_f_ is reached (also called the “forgetting rate”), at which point the distribution of the recalled response is completely uniform. 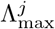 is the maximum recalled spike count of the uniform distribution, taken to be 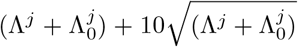 for the rest of the study. We use a value of 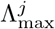 to be 10 standard deviations away from the mean of the largest possible response a cell *j* can have. Though the value of Λ_max_ is arbitrary, the chosen value allows for the uniform distribution in the mixture of recalled responses to cover most of the likely responses generated by the original Poisson distribution. An example distribution of the recalled response is shown in Figure 1d.

In order to deal with a large number of adaptive neurons, we separate them into *P* homogeneous populations of identical neurons. Each population of neurons share the same values of Λ^*j*^, 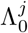, *β*^*j*^, *τ* ^*j*^ and *s*^*j*^. The sum of these i.i.d. Poisson responses is a sufficient statistics of the underlying firing parameter given by Equation (2) as per the Fisher-Neyman factorization theorem [49]. Therefore, the distribution of the sum of responses of a population of identical neurons carries the same information about a sequence **T** as the combined population response across all neurons taken separately. As the sum of *N* i.i.d. Poisson variables is itself Poisson, we can simply multiply the firing parameter of population *j* by the number of cells in said population *N* ^*j*^ to retrieve the correct distribution for the sum of spike responses of a population in Equations (3), (4) and (5). For example, the responses of a network of 500 adaptive cells can be separated into 5 different populations, each of which has 100 identical neurons, as shown in Figure 2a. Each of these populations has a distribution for the sum of its responses shown in Figure 2b. We also take the recall of previous sums of responses to be more uniform as the time since the recalled event increases. From this point onward, we refer to 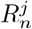 as the sum of the spiking responses of population *j* of size *N* ^*j*^ during event *n* which is described by the redefined firing parameter

**Fig. 2:**
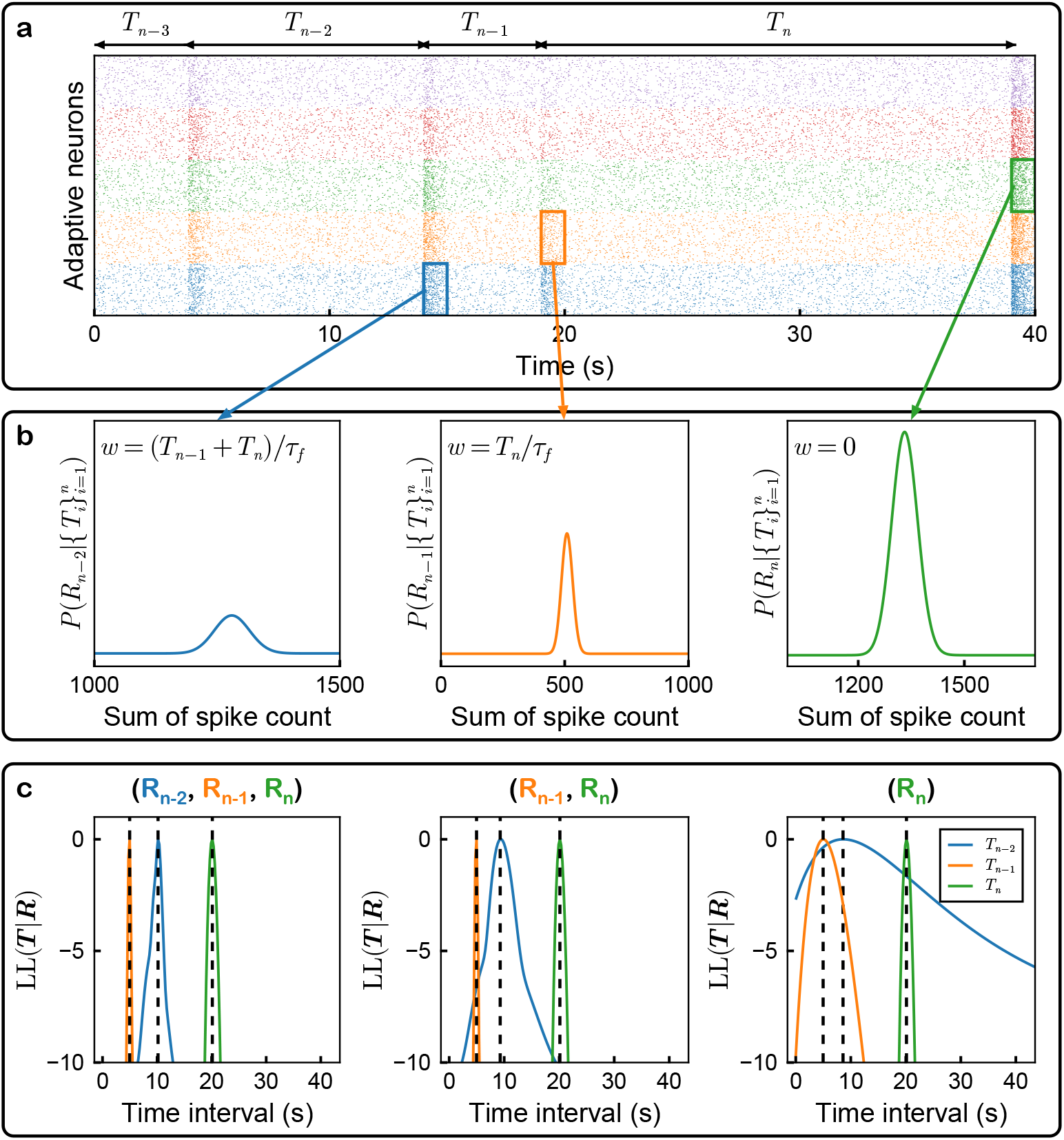
Combining multiple populations to decode sequences of events using maximum likelihood estimation. **(a)** Example spike raster of adaptive neurons with response threshold (RT) of 5 different populations of 100 identical neurons each. Different populations have different colours. Shorter intervals between events yield a fainter response (e.g. event *n* −1) while longer intervals yield a stronger one (e.g. event *n*). A non-zero response threshold (RT) yields baseline-responsive neurons if their resource variable hasn’t recovered enough (e.g. top population during event *n* −1). The populations have threshold values of, from bottom to top, *s* = 0 (blue), *s* = 0.125 (orange), *s* = 0.25 (green), *s* = 0.375 (red) and *s* = 0.5 (purple). **(b)** Distribution of the sum of spikes in a population of 100 identical neurons. The distribution is shown smooth for illustrative purposes; the actual values are discrete. Working memory (WM) is modeled with a mixture between the original Poisson distribution of spike counts and an uninformative uniform distribution, representing a noisy recall of past responses. The weight *w* of the uniform distribution increases linearly with the time elapsed since the recalled event (see Equation (8)). **(c)** Loglikelihood (LL) of time interval values in a sequence **T** given an observed set of responses and recalls **R** from the 5 different populations. The recovery of sequences can be done using maximum likelihood estimation (MLE, see Equation (12)). The dashed lines are the maxima of the LL (i.e. the most likely time interval value) given a realization of responses to the last event and recalls of the responses to previous events. These values were subtracted from the full curve such that the maximum is always at LL=0. This was done to have the y-axis of all 3 panels in a similar range of values for visual purposes. The estimates of past intervals become more accurate (thinner shape) when using more recalls of past responses (increasing from right to left). The top of each panel shows which recalls and responses were used for the MLE. Each loglikelihood curve is plotted by fixing the other time intervals to their real values. The real sequence of events is the same for all panels, namely *T*_*n*−2_ = 10 s, *T*_*n*−1_ = 5 s and *T*_*n*_ = 20 s.

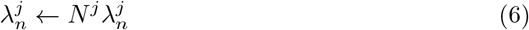

and the redefined maximum recalled spike count

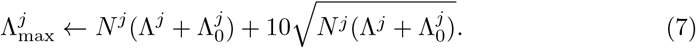

When looking at the responses of multiple populations, forgetting should be happening to the responses of all populations at the same time. Indeed, there is no reason why the previous response of population *j* should be remembered while the response of another population *j*^′^ is forgotten. As such, we write the equivalent of Equation (5) for multiple populations as

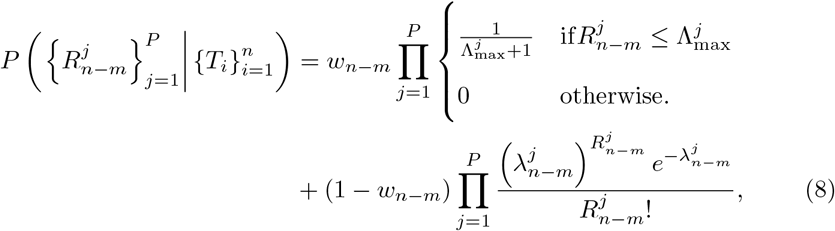

i.e. a mixture of products of independent distributions rather than a product of the independent mixture distributions. This insures that each population cannot be considered as an independent channel in WM that would add unrealistic redundancy as the number of populations is increased, even if they are all identical. Recall is then a network-wide process.

### Bayesian interpretation of working memory

The previous responses are unlikely to be stored in WM as is. Rather, we expect some representation of these responses to be encoded in WM. To justify the use of our mixture model, we adopt a Bayesian perspective. Recall that, as per Bayes’ theorem, the likelihood that the sequence 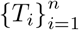 is responsible for the observation of responses from *P* populations 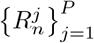 during event *n* is given by

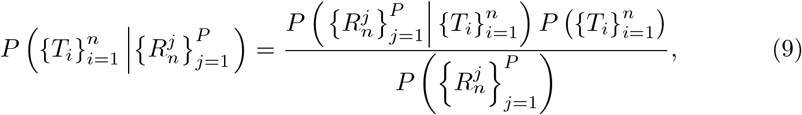

where 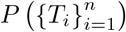 is the prior distribution over the different possible sequences. If no prior information is available, this prior distribution is uniform over all possible sequences. We can then recover the underlying sequence given the responses of adaptive cells using an estimator such as the maximum likelihood estimator (MLE), which simply consists in finding the sequence which maximizes the r.h.s. of Equation (9).

In this work, we consider the realistic strategy where that prior information is indeed available for the estimation of the sequence of events. We therefore replace the prior with

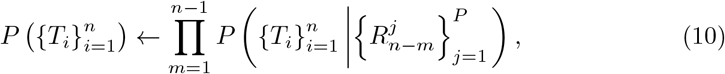

where the product is taken over all of the previous responses during the sequence of events. Equation (10) is what we assume is stored in WM; its exact description would depend on the characteristics of the network that implements WM. To make as few assumptions as possible on the exact implementation of WM, we apply Bayes’ theorem once more on Equation (10):

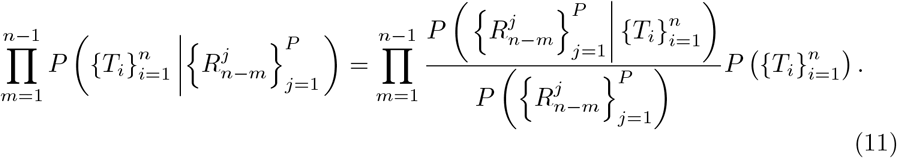

We then use Equation (8) as the likelihood term which can be injected in Equation (9) to obtain the complete posterior distribution of the sequence informed by the prior due to previous responses:

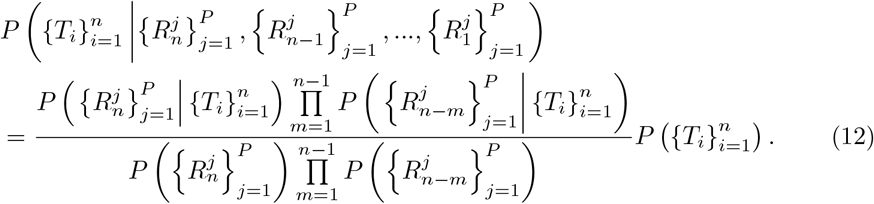

The mixture then describes how well the prior is stored in WM and how its information content degrades over time. Though the above formulation is general and can be implemented with any product over *m*, we choose to have access to all of the *n* − 1 recalls and completely control the capacity of WM through the forgetting rate *τ*_f_. When *τ*_f_ = 0, no WM is available as all recalls are sampled from an uninformative uniform distribution. The presence of WM is therefore controlled by *τ*_f_ *>* 0.

### Arguments for and against using MLE

We could use MLE by maximizing Equation (12) w.r.t. 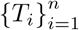 given a sample set of responses and recalls and then retrieve an estimate of the time sequence that generated them. As more recalls of past responses are added to the posterior, the estimates of the time interval values become more precise. An example of this is shown in Figure 2c, where the loglikelihood (LL, the logarithm of Equation (12)) is shown as a function of the time interval values in a sequence of *n* = 3 intervals. We subtract the maximum value of the LL for each curve such that the maximum is always at 0 for illustration purposes. This doesn’t change the location of the maximum. Similarly, the denominator of Equation (12) need not be computed, as it is independent of the sequence 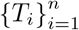. This is one of the advantages of MLE: the costly marginals are not necessary to analyze the model.

The example shown in Figure 2c is one that is somewhat well behaved. Another realization of the estimation process with a different set of responses and recalls could yield likelihoods with substantial irregularities due to the nonlinear nature of the rectification when *s >* 0 and the mixture of the uniform and Poisson distributions. Such an example is shown in Supplementary Figure 1. In that example, the inclusion of **R**_*n*−2_ makes the solution given by MLE drift away from the real value of *T*_*n*−2_ (towards a larger value in this case), indicating a possible bias in the MLE. It is worth noting that the LL curves in Figure 2c are plotted while keeping the other time intervals at their real values. This is not realistic, as MLE would be performed in the 3 dimensions of sequence space (*T*_*n*_, *T*_*n*−1_, *T*_*n*−2_). Nevertheless, it shows how, for example, the confidence on *T*_*n*−1_ and *T*_*n*−2_ is increases when noisy recalls of *R*_*n*−1_ and *R*_*n*−2_ are added to the likelihood by narrowing the LL around the maximum.

Using MLE to retrieve sequences from responses and using those estimates to generate an optimal distribution of adaptation parameters comes with many numerical challenges. One strategy could be to stochastically compute the average error of the estimates made with MLE and minimize this quantity over all the possible sequences. A more analytical approach would be to use the Fisher information (FI), which is given by the following expression for independent adaptive neurons:

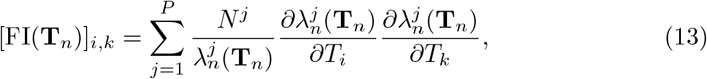

where *i, k* = 1, 2, …, *n*. Equation (13) is a matrix that measures the local — in the *n*-dimensional space of sequences **T**_*n*_ — invertibility of the MLE, i.e. how well can **T**_*n*_ = (*T*_1_, *T*_2_, …, *T*_*n*_) be inferred by the responses 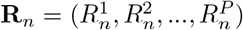, on average.

Note that Equation (13) is for the problem without WM, but including WM would yield a similar expression. Recalls of past responses can be considered as separate populations of noisy adaptive neurons, and their contribution to the FI is additive. FI can be useful as it is related to the lower bound of the error made on the estimates of **T**_*n*_, called the Cramér-Rao lower bound (CRLB):

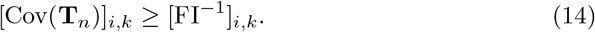

This bound is reached by the MLE when there is a large enough number of neurons, as a small *N* yields a significant bias in the estimates [46]. For smaller *N*, another estimator would need to be designed to reach performance close to the CRLB. Assuming such an estimator exists, a strategy to retrieve the optimal adaptation parameters could be to find which ensemble of parameters minimizes the CRLB, though which quantity to be minimized could be debated. For example, should the sum of the diagonal of [Cov(**T**_*n*_)]_*i,k*_ be minimized, or should its determinant? Moreover, Equation (13) is not appropriate for populations with a RT. Indeed, the FI for a sequence **T**_*n*_ such that *x*_*n*_(**T**_*n*_) *< s* is identically 0 (assuming Λ_0_ *>* 0) since *∂λ*_*n*_*/∂T*_*i*_ = 0. This implies that this population does not contribute to the encoding of this particular sequence. However, it is quite obvious that a population that remains silent does carry information about the sequence. Indeed, it implies that **T**_*n*_ is most likely in an area of sequence space that does not generate a response of the thresholded population. This is not represented in FI.

### Mutual information instead of MLE

Due to these challenges, we instead analyze the mutual information (MI) between the sequences of time intervals and the responses and recalls of populations of adaptive cells. To simplify the notation, we define the vector 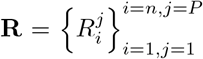 as the set of responses and recalls of the *P* populations and the vector 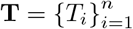 as the sequence of encounters of length *n*. We use the complete set of recalls of past responses (*m* goes from 1 to *n* − 1 in Equation (12)) and the latest responses and control the presence of WM through the value of the forgetting rate *τ*_f_ : *τ*_f_ = 0 is the no WM case, while *τ*_f_ *>* 0 is the WM case. The subscripts of **R** and **T** and the prime superscript of **T** represent different instances of those sets (e.g. different samples of those vector quantities). The MI can be generally written as:

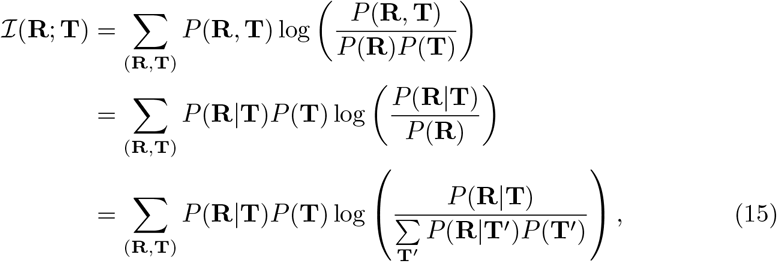

where the sums are done over all possible values of (**R, T**) and **T**^′^. From our model definition, a larger response is usually associated with a longer time interval. This simple fact can help understand what is being calculated in Equation (15). Suppose that we have one population without WM and a sequence of length *n* = 1. In this example, both the response *R* and the interval *T* are scalar values, hence the lack of boldface symbols. Looking at the contribution of one specific pair of response and interval (*R, T*), we immediately see that if *R* is large while *T* is small, then the weight of the contribution of this pair given by the prefactor *P* (*R, T*) = *P* (*R*|*T*)*P* (*T*) is small. Conversely, if both *R* and *T* are correspondingly large, then so is the contribution of this pair to the MI, since their weight *P* (*R, T*) is more important. The fraction inside the logarithm compares how likely it is that an interval *T* generates response *R* to how often the response *R* is observed, regardless of the interval.

To have manageable computation time, we limit ourselves to sequences of up to 3 time intervals. Moreover, we discretize the possible values of time intervals on a finite grid. These simplifications allow us to estimate the MI between responses and sequences and its derivatives w.r.t. the parameters *β, τ, s, N*, Λ and Λ_0_ (see section Methods for a full description). The discretization of the time interval values also means that there is a theoretical maximum of the MI. For a sequence of *n* independent intervals, this maximum is given by the entropy of the time interval sequence distribution

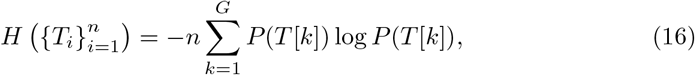

where *G* is the size of the grid (30 here) and *T* [*k*] is the value of the interval at position *k* on the grid. Note that we use different values of *T* [*k*] to represent different distributions, each being the edge of equiprobable ranges on the analyzed support of time intervals. As such, Equation (16) is always given by *n* log *G*. Unless otherwise stated, a uniform distribution of time interval values between 1 s and 30 s is used here.

### Heterogeneous adaptation maximizes information content about sequences

Increasing the total number of adaptive neurons *N*_total_ progressively increases MI towards this limit, as can be seen in Figure 3a. It was previously shown that at least *n* populations of different adaptive neurons (without WM) are necessary to encode a sequence of length *n* [46]. As such, we separated the neurons into *P* = *n* populations whose parameters *β* and *τ* (*s* = 0 for all populations) were optimized through gradient ascent (see section Methods). In both the *n* = 2 and *n* = 3 cases, having WM allows for the network to reach the saturation level of MI sooner in terms of total number of neurons.

**Fig. 3:**
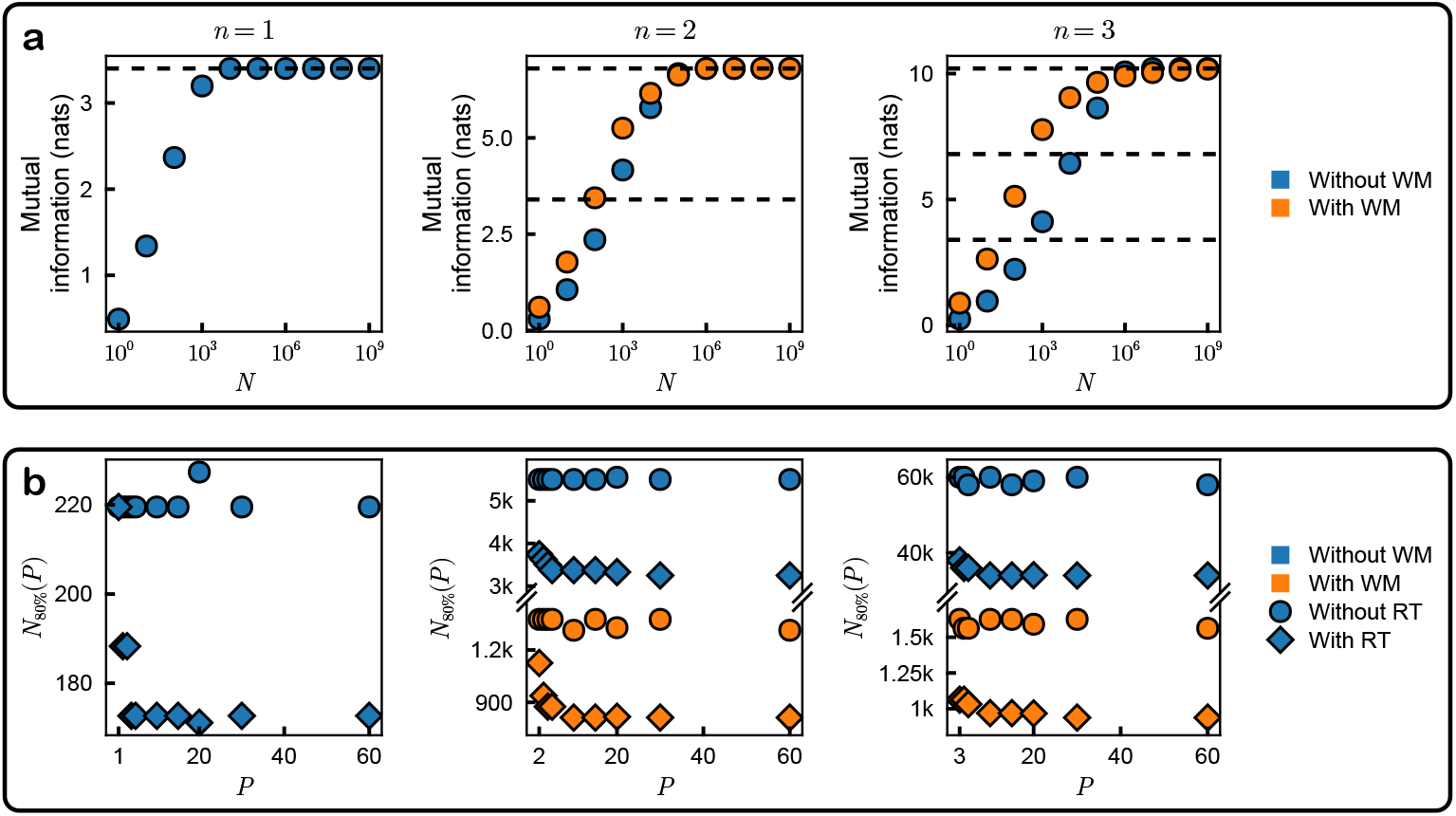
Parametric heterogeneity encodes time sequence information more efficiently. **(a)** Mutual information (MI) content about sequences of different lengths as the total number of neurons N is increased for different numbers of optimized populations (*P* = *n*). The dashed lines are the maximum MI values for sequences of *n* = 1, 2, 3 intervals (see Equation (16)). The populations do not have any response threshold (RT). Including working memory (WM) allows for significantly faster saturation. **(b)** Total number of neurons *N*_80%_ necessary to reach 80% of the theoretical maximum of MI (see Equation (16)) as the number of optimized populations *P* is increased. Including either heterogeneous RT or WM reduces the value of *N*_80%_ significantly. For all sequence sizes, increasing the number of populations only has an effect when RT is available. When WM is available, *τ*_f_ = 35, 65 s for *n* = 2, 3, respectively.

To quantify how much parametric heterogeneity affects MI content, we optimized networks with different numbers of populations with gradient ascent and looked at how many total neurons are necessary to reach 80% of the theoretical limit given by Equation (16), hereby called *N*_80%_. These *N*_80%_ neurons are separated into *P* populations such that 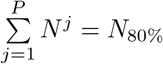. During optimization, we keep the total number of neurons *N*_total_ constant for an ascent step. After the step, we scale *N*_total_ so that it is close to *N*_80%_ (see section Methods). As such, the objective that is optimized is one of efficiency rather than absolute MI: an increase in MI at constant *N*_total_ during a gradient ascent step reduces *N*_80%_. We show the behaviour of *N*_80%_ for optimized networks at different population count *P* in Figure 3b. When there is no RT, we observe no difference as *P* increases. However, allowing for heterogeneous RT increases performance significantly, allowing for fewer neurons to have the same amount of information about the sequences. Incremental efficiency increase can also be observed as *P* is increased. The use of WM also decreases *N*_80%_ significantly. Similarly to the no WM case, increasing *P* only has an effect when RT is available. However, this effect is relatively less pronounced than in the case without WM.

The previous results can be understood quite clearly once we look at the distribution of the optimized parameters of a network with 60 populations (see Figure 4). Indeed, when no RT is available, all of the optimized populations are lumped into *n* or fewer effective populations (clusters) of identical parameters (panels **(i)** and **(iii)**). Once RT is available, however, the distribution of *τ* and *s* become more spread out in each cluster while the values of *β* remain unchanged. One cluster always has *β* = 0 (non-history cells), while the others have values near *β* = 0.5 (history cells). RT thus makes it more efficient to increase the number of different populations, which is what we see in Figure 3b. The effect WM has on those clusters is to change their locations and relative weights. Increasing the value of *τ*_f_ makes the distribution of parameters more similar to the distributions of lower *n* by increasing the proportion of the cluster of non-history cells, as shown in Figure 4iii-iv. There is therefore a trade-off between using the WM to encode previous responses and the built-in history dependence of their responses through *β*.

**Fig. 4:**
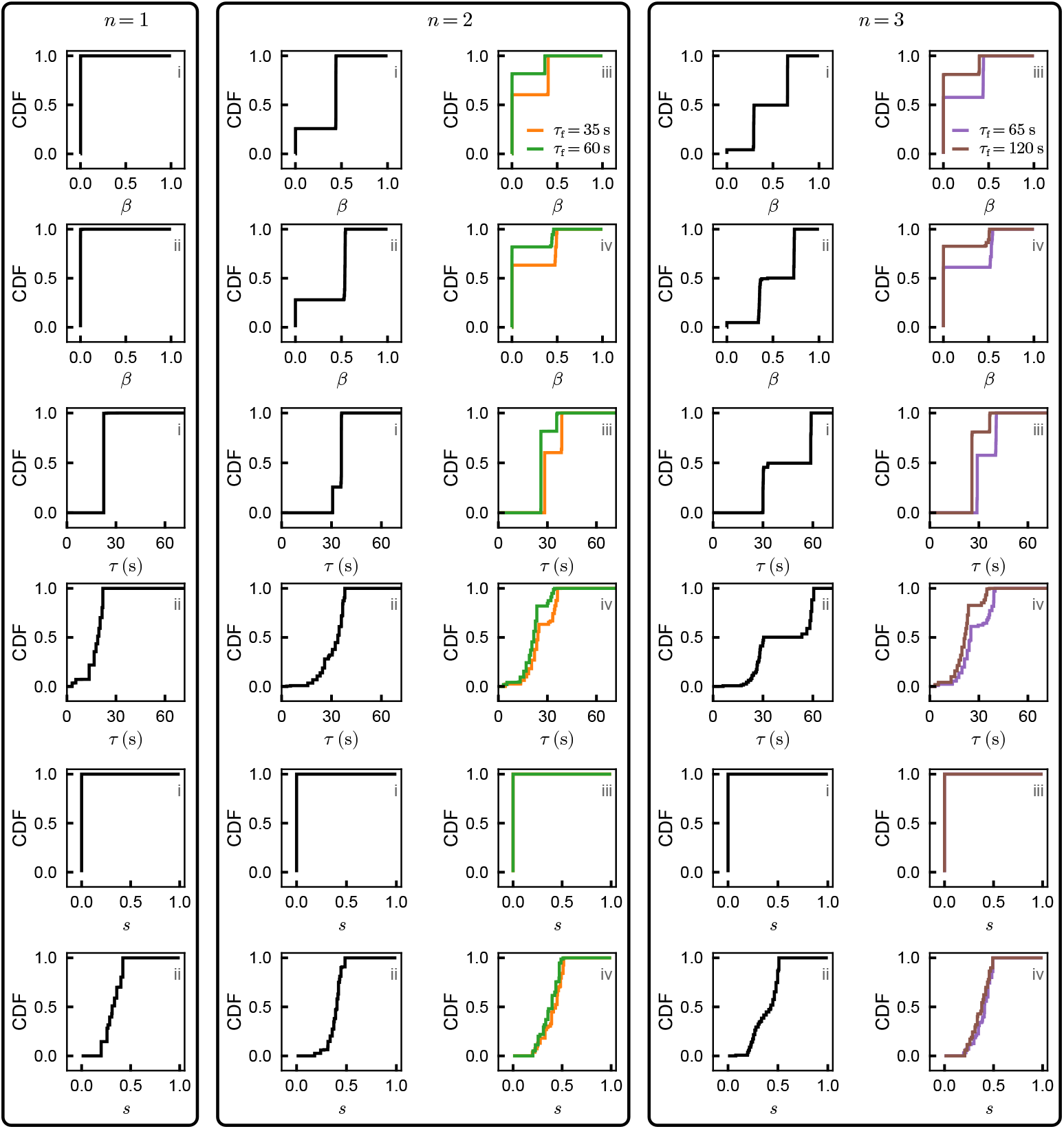
Response threshold heterogeneity enables a continuum of parameters within clusters of effective populations. Cumulative density function (CDF) of the parameters of 60 populations optimized to encode different sequence lengths *n*. **(i)** Without response threshold (RT) and without working memory (WM). The optimization lumps the 60 initial populations into *n* effective different populations (clusters), thus demonstrating the minimal necessary heterogeneity. **(ii)** With RT and without WM. While the history dependence parameters *β* are still lumped into *n* values, the populations now have a continuum of values of *s* and associated *τ*. **(iii)** Without RT and with WM. The populations are lumped in *n* or fewer effective identical populations. **(iv)** With RT and with WM. Neurons in each cluster have identical values of *β* and a continuum of values for *τ* and *s*. For the case *n* = 3, including WM makes the distribution more similar to the *n* = 2 case. For the *n* = 2, 3 cases, increasing *τ*_f_ makes the distributions more similar to the *n* = 1 case, i.e. the proportion of *β* = 0 neurons becomes larger. The use of *τ*_f_ = 35, 65 for the cases *n* = 2, 3, respectively, was done to have a WM that is not overly performant while keeping the complete prior of time interval values in the recalls of responses. This insures that intervals *T*_1_ = 30 s for *n* = 2 and *T*_1_ + *T*_2_ = 60 s for *n* = 3 do not automatically mean that the recall of past responses fails as would be the case if *τ*_f_ ≤ 30 s for *n* = 2 or *τ*_f_ ≤ 60 s for *n* = 3. Legends are shared column-wise.

The idea of clusters of populations becomes much clearer when looking at the marginal distribution of each pair of parameters shown in Figure 5. Indeed, the initial point clusters in the case without RT (panels **(i)** and **(iii)**) remain approximately at the same values of *β*, while the values of *τ* and *s* become more spread out in the case with RT (panels **(ii)** and **(iv)**). These marginal distributions also show a clear quasi-linear relationship between the values of *τ* and the values of *s* within these clusters. This is expected, as *s >* 0 yields neurons that respond only on a given range of time interval values where a different value of *τ* optimally encodes those intervals.

**Fig. 5:**
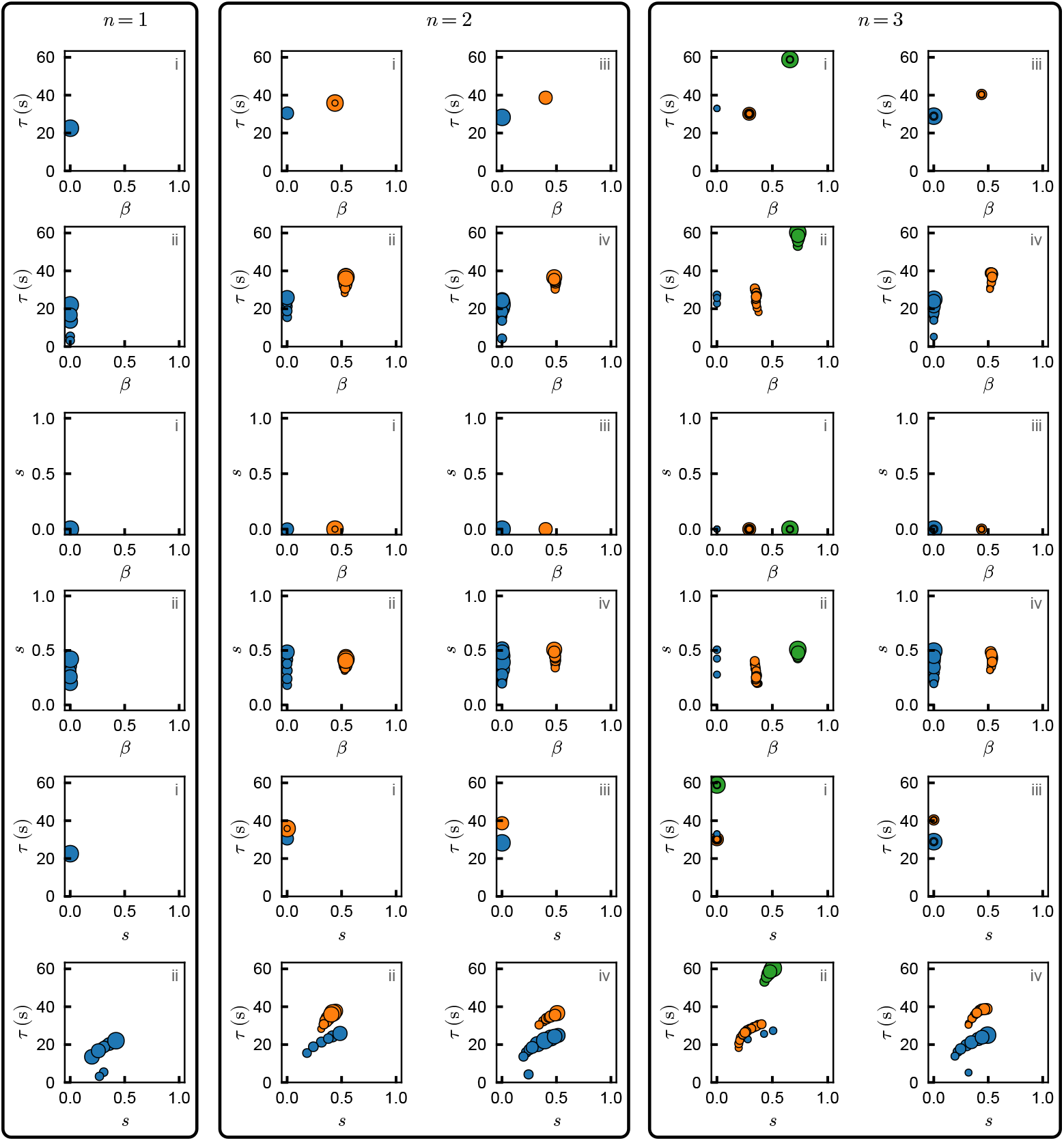
Marginal distributions of parameters show persistent clusters. Marginal distributions of each pairs of parameters optimized for sequences up to *n* = 3 intervals. A total of *P* = 60 populations were optimized and they are each represented by a marker whose size is proportional to the number of neurons they contain. The populations are kept only if they contribute more than 1% of the total neuron count in the network. **(i)** Without response threshold (RT) and without working memory (WM). Parameters are separated into *n* different clusters. **(ii)** With RT and without WM. The values of *β* are still separated into *n* clusters, but the values of *τ* and *s* within these clusters are continuously distributed. **(iii)** Without RT and with WM. The location of the clusters change slightly compared to case in panel **(i)**. In the case *n* = 3, the number of clusters is reduced from 3 to 2. **(iv)** With RT and with WM. The values of *β* are still separated into *n* or fewer clusters with a continuous distribution of *τ* and *s* whose location is slightly different from the case in panel **(ii)**. The colour of each population is associated with its value of *β*. In panels **(iii)** and **(iv)**, *τ*_f_ = 35, 65 for *n* = 2, 3, respectively.

### Heterogeneity tiles the stimulus prior

To better understand how heterogeneity affects the encoding of the values of the time intervals in a sequence, we decompose the MI into stimulus-specific values. To do so, we introduce two measures: the specific information (SI) and the conditional specific information (CSI). While the SI allows to calculate the information content of a specific time sequence, the CSI can do so while completely decoupling the interaction between the different intervals in the same sequence. As such, information content can be visualized as a function of a 1-dimensional variable *T*_*k*_, the value of the *k*-th interval in a sequence. We first use the definition of the SI of a particular sequence **T** = **t** contained in responses and recalls **R** given by

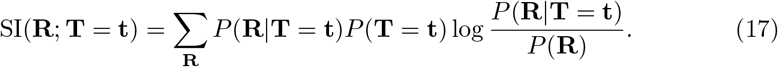

This quantity is also called the “stimulus specific surprise” [50] and is the only positive decomposition of MI [51]. It sums to the MI, i.e.

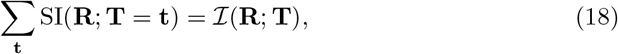

and it compares the likelihood that the responses and recalls **R** were generated by the sequence **T** = **t** (numerator of Equation (17)) to the probability of observing **R** regardless of the sequence that generated it (the denominator of Equation (17)). Looking at one term of the sum over **R**, we see that if both quantities are identical, then it becomes impossible to associate **R** to the sequence **T** = **t** and the contribution of this pair (**R, T** = **t**) to the SI is zero. Conversely, if **R** is generally unlikely (i.e. *P* (**R**) is small) while **T** = **t** often yields such **R** (i.e. *P* (**R**|**T** = **t**) is large), then the ratio is large and so is the contribution of this pair (**R, T** = **t**) to the SI. In other words, observing an unlikely (surprising) response is much more informative about this specific sequence than observing a common one.

To view the effect of allowing heterogeneous RT, we can take the simplest case where *n* = 1 and look at its SI as shown in Figure 6. We reused the optimized parameters of the 5 populations for which we calculated *N*_80%_ in Figure 3b for the case *n* = 1 without RT (*s* = 0, circles in Figure 3b) and with RT (*s >* 0, diamonds in Figure 3b). Each coloured curve is the SI of one of those 5 populations and the thicker black line is the SI of all 5 combined. When *s* = 0 (left), most populations are suppressed into having a very low cell count while only one population (hidden behind the full SI line) remains and encodes all of the possible values of time intervals. Conversely, when *s >* 0 (right), each population’s SI is spread across time interval values in a different way. One population (purple) responds to all intervals, though the SI quickly drops as *T*_1_ increases. The other 4 all have the same behaviour: constant SI until some value of *T*_1_ is reached, at which point the SI jumps up to then follow a similar wide u-shape path. Note that, although those populations do not respond above baseline until their respective threshold is reached, their SI value is still non-zero under said threshold. This constant SI indicates that non-activated cells still carry information: the interval is likely to be a value under the threshold.

**Fig. 6:**
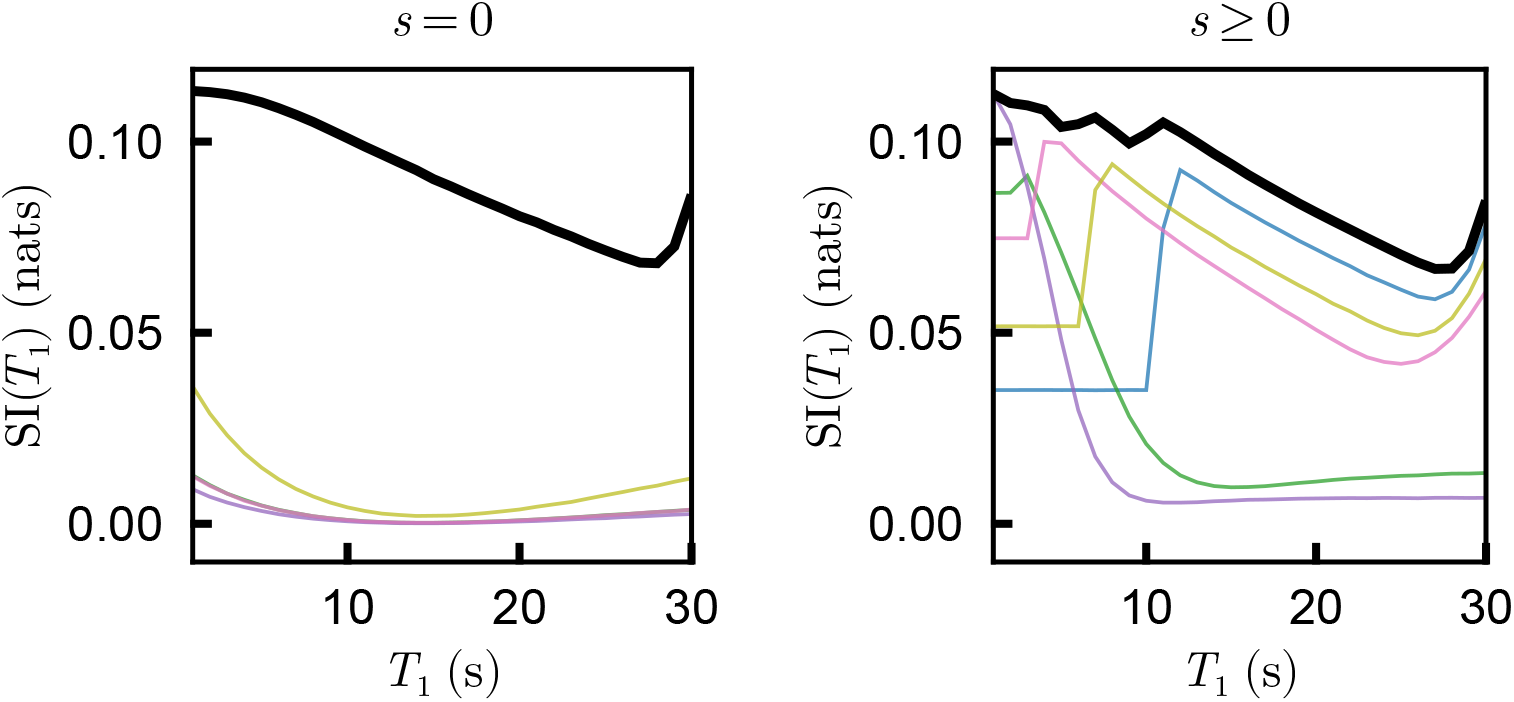
Response threshold heterogeneity allows for a distributed encoding of time intervals. Specific information (SI, see Equation (17)) between responses of individual populations and a single time interval value (*n* = 1). Each colour represents the SI of one population. The parameters of the 5 populations are the ones that were obtained in Figure 3b. The thick black line is the SI of the whole network. When no response threshold (RT) is available (*s* = 0), most populations are suppressed and they therefore have a low SI while only one carries most of the information content, which is behind the line representing SI of the whole network. When RT is available (*s >* 0), populations become specialized to encode specific ranges of time interval values.

Since the SI in Equation (17) is *n*-dimensional, it becomes more difficult to visualize its behaviour as **T** = **t** changes when *n >* 1. To have a metric for a particular interval *T*_*k*_ in a complete sequence **T** = (*T*_1_, *T*_2_, …, *T*_*n*_), we could evaluate the SI when *T*_*k*_ = *t*, while the others are marginalized (i.e. averaged over) as such:

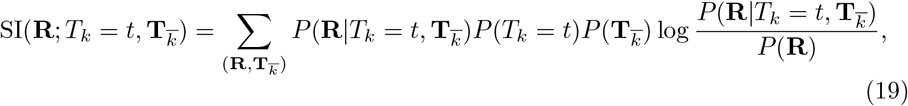

where we separate the complete sequence **T** of length *n* into a single interval *T*_*k*_ and its complementary set 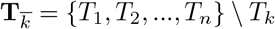.

One problem with this metric is that it is tainted by the information content about the other intervals in a sequence and we can observe the consequence of this with a simple example. Assume that we have a population of non-history cells (*β* = 0) that encodes a sequence of length *n* = 2 with no WM. Because its response does not depend on *T*_1_, the information content about *T*_1_ in the response of this population is null. However, writing Equation (19) for this case, we get

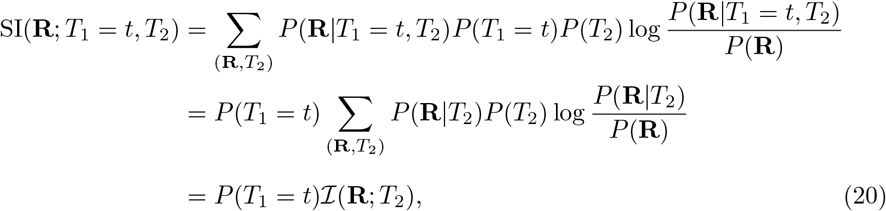

i.e. a non-zero value that only depends on the prior distribution of *T*_1_. For our uniform grid, this value is constant. It would be much more useful for the information metric about *T*_1_ to be 0 in this specific case, otherwise this could be confounded with the behaviour of neurons with a RT where SI remains constant when *x < s* as shown in Figure 6.

For this purpose, we imagine that an oracle is available and knows all of the values of the other intervals in the sequence 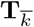. With the help of this oracle, we can compute how much information the responses and recalls contain about only *T*_*k*_ using the CSI:

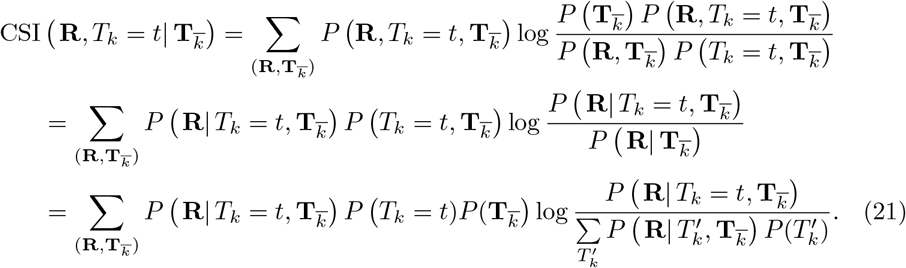

The expectation is taken over the values of **R** and 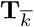. We interpret this quantity as quantifying how much information about *T*_*k*_ = *t* is contained in **R** given that we know the values of the other intervals in the sequence 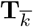 with the help of the oracle. The fraction in the logarithm of Equation (21) compares the likelihood that **R** was generated by a sequence where *T*_*k*_ = *t* and 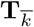 to the probability of observing the same **R** given 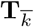 regardless of *T*_*k*_ = *t*. A response that is rare given 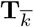 (small denominator), but often generated when *T*_*k*_ = *t* (large numerator) will carry a large amount of information about this specific value of *T*_*k*_ = *t*. Conversely, if the responses or recalls carry absolutely no information about *T*_*k*_ — *T*_*k*_ has no effect on the value of *R* — then this quantity will be identically 0, as intended. The only difference between the SI and the CSI is that the latter uses the probability of observing **R** conditioned on the other intervals 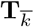 instead of the complete marginal probability of observing **R** in the denominator of the logarithm.

For example, if we again have a sequence of length *n* = 2 and a single population of non-history cells without WM, then we have

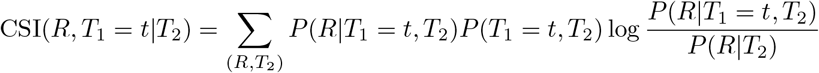

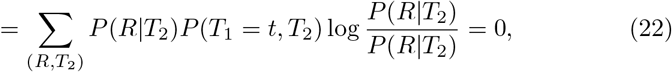

since the response is independent of *T*_1_ when *β* = 0. Similarly, for the same population, we have

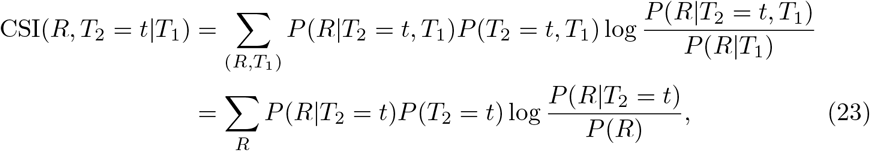

as *P* (*R*|*T*_1_) = *P* (*R*) and *P* (*R*|*T*_2_ = *t, T*_1_) = *P* (*R*|*T*_2_ = *t*).

The CSI allows us to separate the information content about a particular interval

*T*_*k*_ from the rest of the sequence 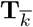. Having a fixed value of *t* also allows us to sweep over the possible values of time intervals, giving an idea of how well the adaptive response encodes a particular interval value.

Note that this represents the ideal case where an oracle with perfect prediction ability of the other intervals 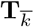 is available. As such, CSI should be considered an upper bound of the actual information about a specific value of *T*_*k*_, because, in reality, an error in the estimate of *T*_*k*′≠*k*_ can influence the error in the estimate of *T*_*k*_. However, as the network’s MI content is optimized towards the saturation bound *H*(**T**), the CSI becomes more and more representative of the MI content about a specific interval.

We compute the CSI for all sequence lengths with the parameters of the 30 previously optimized populations for Figure 3b and show their value in Figure 7. Note that the SI and the CSI are the same quantity in the case *n* = 1. Without RT (*s* = 0, panels **(i)** and **(iii)**), the clusters of populations contribute to the CSI in a similar manner: more information is contained at the extrema of *T*, though their exact shape is different depending on which cluster a population is in. With RT (*s >* 0, panels **(ii)** and **(iv)**), the CSI curves now have a maximum value of *T* which is different from population to population. In the cases *n >* 1, we can also observe that some populations have no information about *T*_*n*−1_ and *T*_*n*−2_ when there is no WM (panels **(ii)**). These are the populations of non-history cells (*β* = 0). Moreover, the threshold appears to only affect the last interval and not the previous ones when there is no WM (panels **(ii)**). When WM is available, the thresholds affect all intervals, though the CSI is progressively lower for past intervals (panels **(iv)**).

**Fig. 7:**
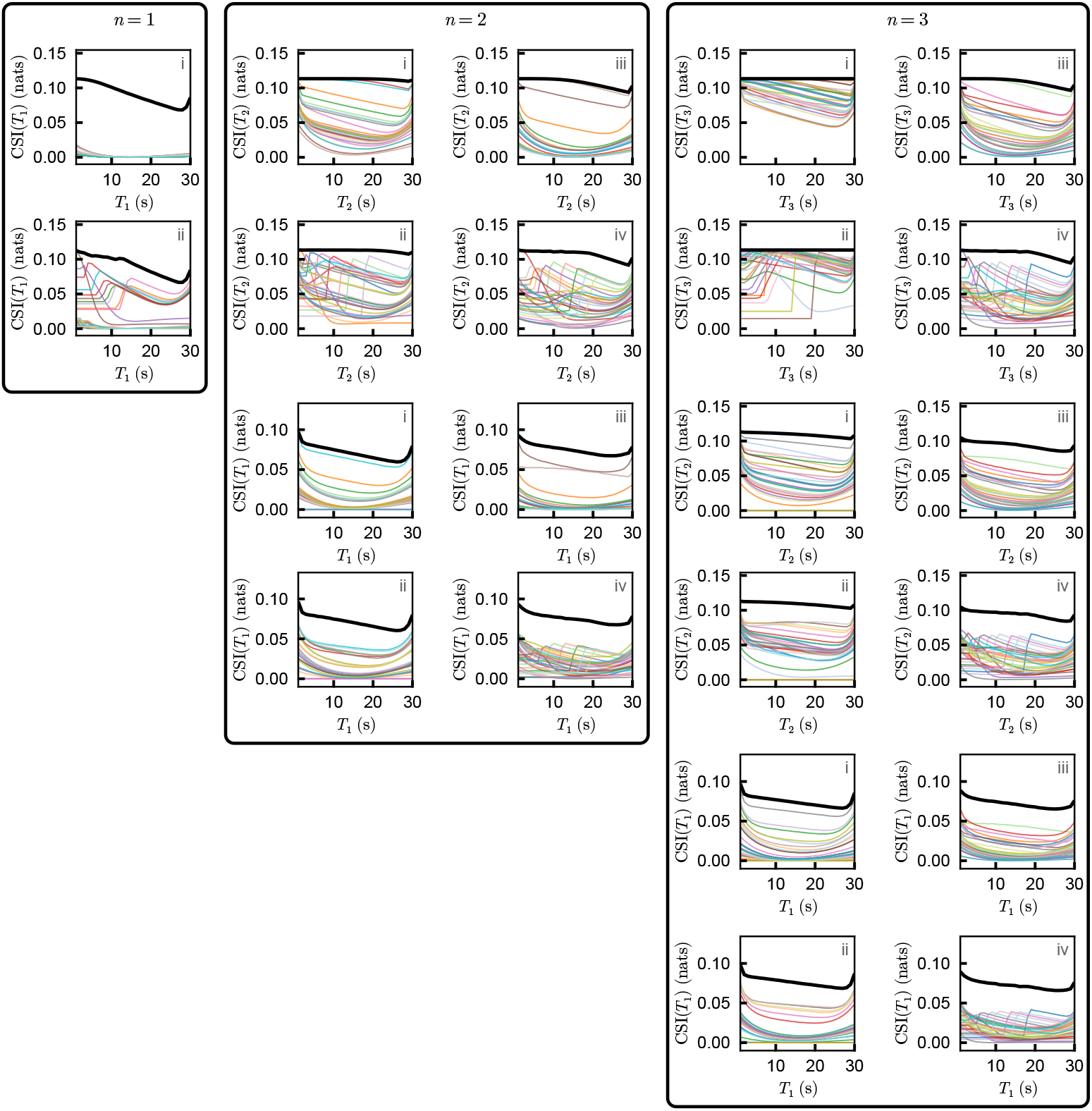
Response threshold helps the encoding of longer sequences with and without working memory. Conditional specific information (CSI, see Equation (21)) between responses of individual populations and individual time interval values in a sequence of *n* intervals. Each colour represents the CSI of one population. The parameters of the 30 populations were optimized simultaneously to maximize mutual information content about the complete sequence of time intervals. They are the same optimized populations as in Figure 3b. The thick black line is the CSI of the whole network. **(i)** Without response threshold (RT) and without working memory (WM). Populations coalesce into *n* clusters of effective populations. Each population has a different weight due to its respective number of neurons. The case *n* = 1 has mainly one population contributing to the CSI which cannot be seen as it is hidden by the total CSI line. **(ii)** With RT and without WM. Populations separate into parameter space such that each one has a specific value of the latest time interval value *T*_*n*_ that is maximally encoded. Previous intervals are encoded similarly to the case without RT. A large portion of the populations does not contain any information about the previous intervals because *β* = 0. **(iii)** Without RT and with WM. Populations coalesce into *n* or less effective populations. All populations encode all intervals in a sequence even when *β* = 0 because of the WM. **(iv)** With RT and with WM. Populations separate into parameter space where each one has specific values of latest and previous time intervals that are maximally encoded. In all cases, the number of total neurons was kept at *N*_80%_. **(ii)** and **(iv)** have a significantly lower number of neurons compared to **(i)** and **(iii)**, respectively. In panels **(iii)** and **(iv)**, *τ*_f_ = 35, 65 for *n* = 2, 3, respectively.

The sudden increase in CSI at a specific value of *T* can be explained by looking at the case *n* = 1. Since the optimal populations are composed of non-history cells, we can write, based on the exponential dynamics of adaptation, the minimum time interval value that will elicit a response from population *j* as

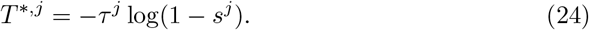

To further analyze the phenomenon, we initialize 30 populations randomly and then maximize the MI about a sequence of *n* = 1 interval 10 times to retrieve a total of 300 optimal populations. We then compute and pool the values of *T* ^*^ from the optimized values of *s* and *τ* for each of the resulting populations and weight them by the number of neurons they have. When we look at the resulting distribution of *T* ^∗^ (see Figure 8a), we see that the values of *T* ^∗^ settle at the values of the grid of time interval values: the thresholds tile the prior distribution.

**Fig. 8:**
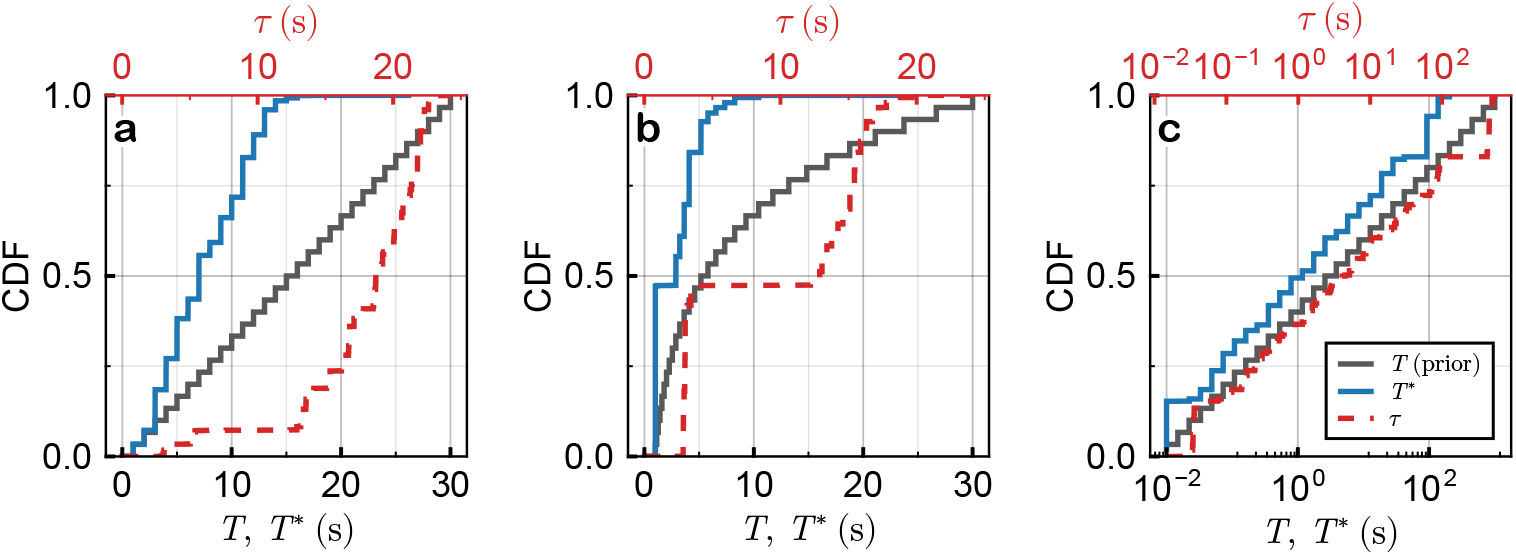
The time thresholds of each population is closely related to the prior distribution of time interval values. Cumulative distribution function (CDF) of the minimum time interval values *T* ^*^ (bottom axis) eliciting a response and the recovery time constants *τ* (top axis) compared to the prior distribution of time intervals (i.e. stimuli, bottom axis). The distributions are constructed from the values of 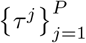 and 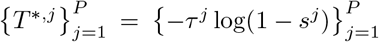 resulting from the optimization of *P* = 30 populations. This is repeated 10 times and all of the resulting 300 values of *T* ^*^ and *τ* are pooled together and weighted by the number of neurons *N* ^*j*^ in their respective population. **(a)** A uniform prior distribution of time intervals yields a linearly increasing CDF of values of *T* ^*^. The associated distribution of *τ* is bimodal, with the second mode also increasing linearly. **(b)** A powerlaw prior distribution with *P* (*T*) ∝ *T* ^−1^ of time intervals yields a high concentration of small *T* ^*^ and *τ* values. Larger values of both variables increase more steadily, with a lower concentration of larger values. **(c)** A scale-free (powerlaw) prior distribution with *P* (*T*) ∝*T* ^−1^ spanning multiple orders of magnitude with time intervals starting from 0.01 s and going to 1000 s. The resulting thresholds and recovery time constants also appear scale-free. The legend is shared by all panels.

We next test whether a different prior distribution decreasing with *T* such as a powerlaw (as seen in some environments) is better tiled by the set of optimized time constants and thresholds. Note that each element *T* [*k*] on the grid of size *G* has the same probability of being drawn given by *P* (*T* [*k*]) = 1*/G*. We simply change the values of *T* [*k*] from being on a uniform grid where *T* [*k* + 1] − *T* [*k*] = (*T*_max_ − *T*_min_)*/*(*G* − 1) to instead being spaced such that *T* [*k* + 1]*/T* [*k*] = (*T*_max_*/T*_min_)^1*/*(*G*−1)^, with *T* [1] = *T*_min_ and *T* [*G*] = *T*_max_. Each element on the grid thus represents the edge value of an equiprobable range between *T*_min_ and *T*_max_ where the continuous equivalent of the distribution of intervals is given by *p*(*T*) ∝ *T* ^−1^.

The resulting distribution of thresholds *T* ^∗^ still sit on the values of the time interval prior (see Figure 8b). In fact, the average deviation from the nearest value of time interval on the defined grid *T* ^∗^ − *T*_nearest_ is 4.3 ms and 1.1 ms for the uniform and powerlaw distributions, respectively. The uniform prior case appears to have a steady increase in the CDF of *T* ^∗^ between the minimum and maximum values of *T* ^∗^, much like the uniform prior CDF of *T*. Something similar can be observed for the powerlaw prior case, where the CDF of *T* ^∗^ is mostly concentrated at small values of *T* ^∗^ and increases more slowly afterwards, much like the powerlaw prior CDF. The close relationship between *τ* and *s* (and therefore *T* ^∗^) means that the CDF of *τ* behaves similarly, though in a bimodal manner. Many cells have an identical small value of *τ* while the others steadily increase in value. We also show the resulting distribution of *T* ^∗^ associated with a powerlaw prior whose support spans multiple orders of magnitude (going from 0.01 s to 1, 000 s) in Figure 8c and see a definite relationship between the prior distribution and the distributions of *T* ^∗^ and *τ*.

We also notice that the values of *T* ^∗^ from MI optimization shown in Figure 8 do not span the complete prior. Rather, the maximum *T* ^∗^ reaches is 14 s and 9 s for the uniform and powerlaw priors between 1 s and 30 s, respectively, while being around 137 s for the powerlaw prior spanning 0.01 s and 1,000 s. This is most likely due to the objective of 0.8*H*(**T**) being reachable more efficiently through the analogue encoding of time interval value with the number of spikes in a response rather than with thresholds. Indeed, if we repeat the simulation of Figure 8, but with an objective of 0.99*H*(**T**), the optimal threshold distribution spans a larger range of *T* ^∗^ values: analogue encoding is no longer enough to reach the target efficiently. This can be seen in Supplementary Figure 3. Note that this much larger objective implies a smaller gradient magnitude which necessitates larger sample sizes (1, 000, 000 samples per gradient estimate here) to have an accurate estimation of the gradient. This is why we limited this higher objective to the *n* = 1 case.

## 3 Discussion

### Summary

Rat hippocampus [42] and weakly electric fish thalamus [45] provide an attractive model to understand how heterogeneity supports the capabilities of neural ensembles to encode and recall time events. Inspired by experimental observations, we use a recovery-from-fatigue adaptation model to recreate observed responses to “encounter” stimulus events. Motivated by the pallial circuitry [52] assumed to decode the electric fish “thalamic” time interval code [45], we also include a simple generic working memory model of previous responses. This allows us to quantify how the information about a sequence of inter-event intervals is distributed across all the responses.

Rather than use a possibly biased Bayesian decoder — such as MLE with a small number of neurons — to retrieve sequences from responses, we turn to the more general quantity of MI. With a few mathematical simplifications, we are able to compute MI and its gradient with reasonable computational time, allowing us to do stochastic gradient ascent on the adaptation parameters, *β, τ* and *s* representing stimulus history dependence (adaptation memory), recovery from fatigue time constant, and single neuron RT, respectively. We encover the effect of heterogeneity in those parameters, including correlations between parameters, by maximizing MI of an increasing number of populations *P* in different situations.

Without threshold heterogeneity, increasing *P* past the sequence length *n* has no significant effect, as the populations coalesce into the same *n* sets of parameters. WM has alters the optimal parameters; but without the flexibility provided by RT, the populations still cluster together. However, when *n* = 3, the number of effective populations changes to 2 (rather than 3 in the no WM case) as the information content about the first interval *T*_1_ due to history cells (*β >* 0) is too low, and therefore taken care of by the WM. RT heterogeneity allows for a richer parameter distribution, where clusters each encode specific time intervals through the specialized selection of thresholds *s >* 0. Accordingly, the associated time constants also become distributed to optimally encode these interval values.

### Working memory

Two types of memory are used here. “History dependence” results from the channel biophysics that carry a fraction *β* of resources after an event. This creates a dependence of the response on the complete sequence of time intervals, rather than on only the latest one when *β* = 0. This separates the types of adaptive neurons into “history cells” (*β >* 0) and “non-history cells” (*β* = 0). The information about past intervals in history cells quickly decays as the total time of the encoded sequence increases. The second type of memory is a simple WM: recalls of past responses mix the original response distribution and an uninformative or “noisy” uniform one whose weight increases as *T/τ*_f_ until it reaches 1. This assumes that the circuit receiving and subsequently remembering input from adaptive neurons is also involved in other computations; storage resources grow more scarce as sequence length and accumulated noise grow, much like in visual WM models [53–55].

The chosen WM model is computationally simple, and we expect qualitatively similar results for different WM models, so long as the quality of recalls deteriorates monotonically with time. Maximizing MI unveiled a trade-off between both types of memory. A smaller *τ*_f_ leads to a weaker WM, making history cells more important to encode sequences. Conversely, a larger *τ*_f_ makes WM more reliable, allowing for fewer history cells. This can be seen in panels **(ii)** and **(iv)** of Figure 4: larger *τ*_f_ increases the proportion of *β* = 0 neurons.

### Heterogeneity in natural and artificial neural networks

Our findings also show that each region in (*β, τ, s*) space has its own purpose for interval coding. The history dependence *β* increases the amount of memory about past intervals available in a neural response. The optimized populations consistently separate into clusters of identical values of *β* with and without WM. One cluster is always made of non-history cells, making it dependent only on the latest interval and thus maximizing the information content about it. When a sequence has more than one interval, the other clusters are made of history cells, each increasing the information content about the penultimate interval and the previous ones.

The decay time *τ* is closely related to *β* and *s*. When *s* = 0 for all cells, *τ* takes the same number of values as *β*, i.e. the optimal values of *β* have their associated optimal values of *τ*. When heterogeneity in RT is allowed, the optimal values of *β* remain lumped together. Within these clusters, however, *τ* and *s* are distributed continuously. This allows for a better shared workload among populations and an increase in efficiency as can be seen in panels **(ii)** and **(iv)** of Figure 7. An abrupt increase in the CSI of a population at a specific value of the time interval *T* ^∗^ (see Equation (24)) indicates that this population was optimized to maximize information content about time intervals beyond this value.

The relation of *τ* to *β* seen in Figure 5 for *n* = 2, 3 reveals that larger adaptation time constants are associated with stronger history dependence in the adaptation process. This makes sense intuitively, as a stronger influence of past events could rely on slower recovery from neural fatigue. Interestingly, this correlation also agrees qualitatively with the one observed experimentally between these parameters [45] (Figure 6 supplement 1), which suggests this could be one of the consequences of a heterogeneity that maximizes input-output MI.

Two levels of heterogeneity appear through MI optimization. The first is the minimum heterogeneity necessary to encode sequences of *n* intervals, which separates the populations into clusters. This is the manifestation of a previously shown result in the same system where the number of different populations needs to grow with the sequence length to enable sequence inference [46] (Figure 8). The second level is the continuous distribution of thresholds and recovery time constants.

This is consistent with literature asserting that neural diversity is computationally desirable rather than simple biological noise. For example, biophysical diversity decorrelates responses of pairs of neurons of the same type [3], increasing the amount of information they carry about a shared stimulus. It enhances discrimination between ethologically relevant sensory stimuli in weakly electric fish [56]. Moreover, moderate heterogeneity — a blend of diversity and similarity — best encodes stimuli [4, 57], resembling how our populations of adaptive neurons separate into clusters of similar cells. At the single-neuron level, diversity has also been shown to support coding across multiple timescales [58, 59] and to promote robust learning in spiking networks [60, 61]. Our results contribute to this literature through a mechanism of division of labour. RT distributions reduce redundancy between populations that become tuned to specific interval ranges while remaining silent beyond them. It remains to be seen how the heterogeneity shaping found here would change if neurons adapted on multiple time scales, e.g. to respond to the many time scales of auditory input [62].

### Coding capacity of silent cells and efficient coding

The efficiency of the uncovered encoding is twofold. First, the optimization yields the best configuration of populations while the total number of neurons is kept near the *N*_80%_ objective (by construction: see Methods), which minimizes the neuron count for a fixed information target rather than maximizes absolute MI. This naturally pushes *N*_80%_ to lower values when possible, increasing the MI per neuron. A second efficiency mechanism emerges from RT heterogeneity: silent neurons. Cells that do not fire during a stimulus reduce metabolic costs. This strategy still enables a silent cell to have information about sequences that do not elicit any responses (e.g. a short interval below *T* ^∗^) as can be observed in panels **(ii)** and **(iv)** of Figure 7. Indeed, these silent neurons still have a non-zero CSI since they “know” the interval is shorter than *T* ^∗^.

Tiling the stimulus prior is advantageous in spite of the loss of dynamic range incurred when the threshold is positive; the informational benefit of silencing outweighs that loss. The tiling seen here conjures up the idea of “histogram equalization”, in which all possible responses are used equally given a stimulus prior [63]. Similar results of threshold distribution were shown in salamander On/Off retinal ganglion cells [64, 65]. The tiling of the prior with thresholds remains in competition with the analogue encoding of intervals using spike counts. For the smaller time interval values, it is better to have populations with a lower RT that code for a wider range of interval values, as can be seen in Figure 8. Similarly, the values of *T* ^∗^ do not span the complete prior, only reaching a value 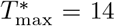 s in the case of a uniform prior. This implies that rate coding of interval duration using spike count is more efficient to encode intervals 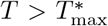 than having populations with thresholds 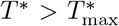 that have a more limited dynamic range.

Silent cells therefore contribute to the encoding of time intervals at reduced energy cost. This is in line with the idea of sparse and distributed coding [66, 67] with highly specialized (often silent) cells observed in hippocampus [68, 69]. Similarly, nonresponsive neurons in the weakly electric fish were shown to contain information about spatial stimuli through their covariance with their responsive counterpart [70].

This silent code decreases the number of neurons necessary to reach the same information level, but also the energy consumption. Indeed, action potentials dominate the brain’s signalling energy budget and energy grows steeply with firing rate [6, 7]. Theoretical work also shows that optimal codes under energy constraints are sparse [71, 72]. Even though we do not consider any metabolic cost of firing rates during the optimization process, the uncovered configuration is still more cost-efficient by having neurons that are quiescent outside their interval band. The heterogeneity we report here can therefore be viewed as both neuron-efficient and rate-efficient, through this sparse code.

### Laplace transform

Our latent variable *x* offers an alternative perspective on the scale-invariant theory of neural time [41, 73], where an ensemble of leaky integrators with powerlaw-distributed decay rates encode the Laplace transform of the stimulus. An inverse Laplace transform read-out of those leaky integrators yields time cell activity similar to that observed in hippocampus. Our adaptation mechanism, where each population has its own time constant *τ* ^*j*^, offers similar coding capacities. The event-based response of adaptive cells offers a snapshot read-out of the recovery process, which can then be decoded for timing information. A similar correspondence has been made for ramping cells [74], and continuous attractor networks can implement such inverse Laplace transforms for decoding [75]. Our model does not map completely onto the Laplace framework. Our latent integrators *x* are updated during discrete events with a depletion factor *β*, which can be mapped to more biophysically detailed cell models [76]. The neural model underlying this Laplace perspective exhibits firing rate adaptation following the onset of a constant input, with the adaptation time constant having different values across the population [44, 77]. After each spike, the voltage is reset, making spiking history irrelevant to this time representation.

The Laplace framework yields Weber-Fechner scaling of inputs [42] that allows the representation of time across multiple orders of magnitudes [78]. Our results show that this type of scaling can be obtained simply through MI maximization (see Figure 8b,c): a scale-free stimulus prior also yields logarithmically compressed recovery time constants, a mathematical representation of the Laplace transform. We also realize that the heterogeneity strategies reported here could be co-opted by other natural and artificial systems to optimally convey temporal information streams beyond event sequences.

## 4 Methods

All numerical simulations were done in the *Julia* programming language. For a sequence of length *n* = 3 with WM and *P* = 60 populations, the running time for the optimization of this network is approximately 4 days on an NVIDIA H100 GPU.

### Estimating the mutual information

We start with the definition of mutual information (MI) as in Equation (15). The main difficulty in calculating the MI lies in the marginal *P* (**R**). Since the sum over **T**^′^ is usually done in a large multi-dimensional space, one can try to estimate the marginal *P* (**R**) using Monte Carlo simulations. Indeed, both the inner and outer sums are expectations, such that a numerical approximation of Equation (15) becomes

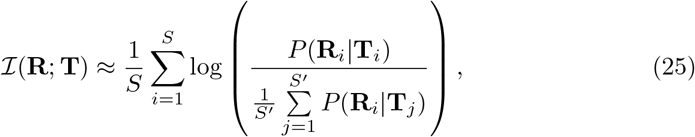

where **T**_*i*_, **T**_*j*_ ~ *P* (**T**) and **R**_*i*_ ~ *P* (**R**|**T**_*i*_). Note that **T**_*i*_ is a vector of size *n* and the subscript *i* indicates one sample of such vector. Similarly, **R**_*i*_ is a vector containing all the responses and recalls of past responses of all populations (size *nP* with WM or size *P* without WM) and the subscript *i* represents a single sample of such vector. *S* and *S*^′^ are the number of samples used to compute the outer and inner expectations, respectively. For each outer sample of **T**_*i*_ and associated **R**_*i*_, we need to sample *S*^′^ new sequences 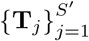 over which we average *P* (**R**_*i*_|**T**_*j*_) to obtain an approximation of *P* (**R**_*i*_). Equation (25) is a nested Monte Carlo (NMC) estimation of the MI, which usually has a convergence rate of 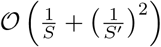. Some empirical laws for the choice of *S* and *S*^′^ can be used to have the lowest possible error given a finite operation budget, though increasing only *S* reduces the variance of the estimator towards a biased value [79]. The convergence rate and the total amount of samples *SS*^′^ necessary for the calculation of Equation (25) quickly becomes astronomically large with the size of the prior stimulus space **T**.

Therefore, a simple way that we can get reasonable compute times is by reducing space of the prior stimulus and by exploiting the highly parallelizeable nature of the model. Since each cell is conditionally independent given a sequence **T**_*j*_, we can compute the values of log *P* (**R**_*i*_|**T**_*j*_) for any **T**_*j*_ given responses and recalls of past responses **R**_*i*_ sampled from *P* (**R**|**T**_*i*_) in parallel. Moreover, the fact that we constrain the possible values of time intervals to a small grid of 30 elements allows us to compute the real marginal given a sample **R**_*i*_. Even with a sequence of length 3, the inner sum needs to be made on 30^3^ = 27, 000 sequence values of **T**_*j*_, which is achievable on modern computer hardware. As such, we propose the following Monte Carlo estimator of the MI:

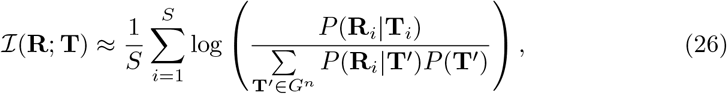

where **T**_*i*_ ~ *P* (**T**), **R**_*i*_ ~ *P* (**R**|**T**_*i*_) and *G* is the chosen finite grid (of size 30 here). We compute each value of the log-conditionals log *P* (**R**_*i*_|**T**^′^) independently and simultaneously for all **R**_*i*_ and **T**^′^. We obtain the denominator of the log of Equation (26) by combining the joint probabilities with

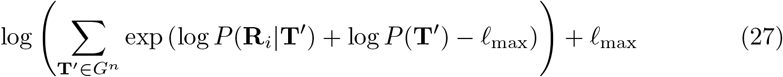

for numerical stability. Here, 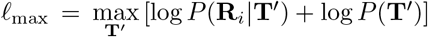 is chosen such that the maximum value in the exponentials is 0, which avoids numerical explosions. Because we compute the true marginal *P* (**R**_*i*_) by summing over all possible sequences **T**^′^, the convergence rate of the unbiased estimator given by Equation (26) is 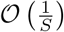.

Because this is a Monte Carlo sum done by sampling a sequence **T**_*i*_ and a corresponding response **R**_*i*_, the contribution weight of each pair (**R**_*i*_, **T**_*i*_) corresponding to the value of *P* (**R**_*i*_, **T**_*i*_) = *P* (**R**_*i*_|**T**_*i*_)*P* (**T**_*i*_) in Equation (15) is done implicitly through the frequency of appearance of said pair in the sum. For example, the response *R* of a single population without WM to a single interval *T* are usually proportional, i.e. a long interval usually produces a large response. As such, an unlikely pair such as a large response *R* to a small interval *T* will occur less often in the sum given by Equation (26) and, as such, the contribution of such pair to the MI will be small. On the other hand, a long interval *T* producing a large response *R* will occur more often, contributing much more to the estimated value of MI.

The computation of the CSI (see Equation (21)) is done in a similar manner, although the expression is slightly different:

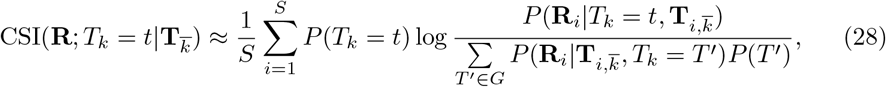

where 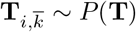 and 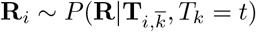.

We proceed in batches when memory is limited until the number of samples reaches *S* = 100, 000, unless otherwise specified. This was done on a graphics processing unit (GPU) using the *CUDA*.*jl* library, which allows for large multithreaded operations.

### Estimates of mutual information gradient

To do gradient ascent on MI, we need to estimate its derivatives w.r.t. each of the population’s parameters. To do so, we first start with the definition of MI in Equation (15) and take its derivative w.r.t. any parameter *θ*^*j*^, which is either *β*^*j*^, *τ* ^*j*^, *s*^*j*^, *N* ^*j*^, Λ^*j*^ or 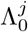. We cannot simply take the derivative of Equation (26), because the sampling of **R**_*i*_ depends on the parameters *θ*^*j*^. We can then get the following expression (see Supplementary Methods for the full derivation):

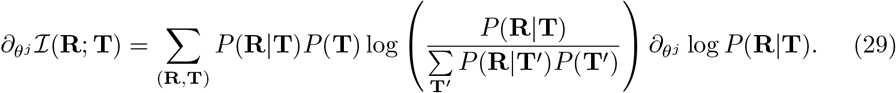

This formulation is sometimes called the score function (SF) [80]. Using the same strategy of doing the explicit inner sum for the marginal distribution over all possible values of **T**, we have the following estimator of the derivative of MI:

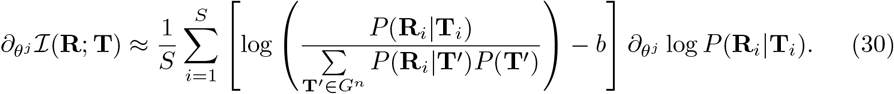

where *b* is a control variate that helps reduce the variance [81], here taken to be the current estimate of MI. Just like the MI estimation, **T**_*i*_ ~ *P* (**T**) and **R**_*i*_ ~ *P* (**R**|**T**_*i*_). This estimator also has the advantage of being highly parallelizable, making it possible to compute log *P* (**R**_*i*_|**T**_*j*_) and *∂*_*θj*_ log *P* (**R**_*i*_|**T**_*j*_) simultaneously for all (*i, j*) and combining them into Equation (30). We again proceed in batches when memory is limited until *S* = 100, 000.

Because we are treating populations of neurons rather than individual neurons, the Poisson distribution of the total response has a parameter 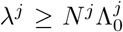. As such, if we have a significant baseline and number of cells in a population, the Poisson distribution can be approximated by a Normal distribution with mean and variance *λ*^*j*^. This approximation allows for a significant improvement of the gradient estimate. Indeed, if we redefine the *i*-th sample of the Normal distribution of the response/recall 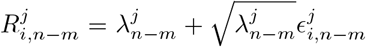, with 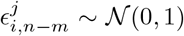, then the derivative can be taken inside the expectation of Equation (26), as we can now compute 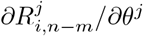 directly. This is because the random sampling of 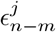 is independent of the parameters *θ*^*j*^. This is often called the reparameterization trick [82] which greatly reduces the variance of such gradients estimators. Note that the uniform distribution in Equation (8) now has a normalization constant of 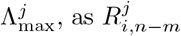 is a continuous variable. A second independent expression for the derivative of the MI can then be written as (see Supplementary Methods for the full derivation)

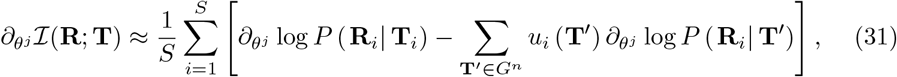

Where 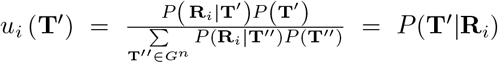 is the posterior probability of sequence **T**^′^ given sample **R**_*i*_ with 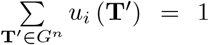. The full expressions for *∂*_*θj*_ log *P* (**R**_*i*_|**T**^′^) for the different parameters of this model are derived in Supplementary Methods.

Supplementary Figure 2 visually compares both gradient estimators by varying one parameter of one population at a time in a network of 3 populations and computing the MI and its derivative w.r.t. the changing parameter for a sequence of *n* = 1 interval. The mean of 10 repetitions of this process each with *S* = 100, 000 (left) or *S* = 1, 000, 000 (right) shows that both estimators indeed estimate the derivative of the MI. The standard error over the 10 repetitions is shown in the shaded areas and shows that, for the same number of samples, the Normal approximation has a significantly reduced variability while keeping the same shape – and thus optima – as the true Poisson estimate. Therefore, the optimization was done using the Normal approximation in the paper.

### Gradient ascent

With these gradient estimators, we do stochastic gradient ascent using the Adam optimizer [83] from the *Optimisers*.*jl* library with a learning rate of 0.01 for all parameters except *s*, which has a smaller learning rate of 0.005 due to it being less stable. Because we have constraints on the domain of the different parameters, the gradient ascent is done on a set of transformed variables that can take any real value, namely:

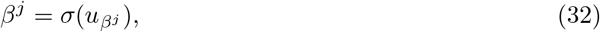

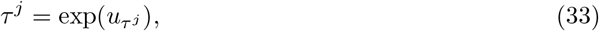

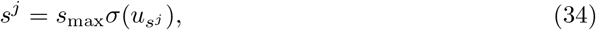

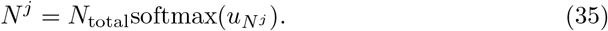

where *σ*(*x*) = 1/(1 + *e* −*x*) and

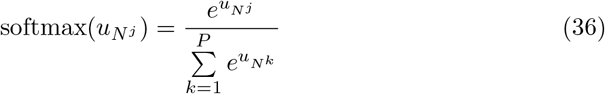

such that 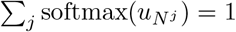 and Σ*j N* ^*j*^ = *N*_total_, always. These definitions allow us to do the optimization in the unconstrained space (from −∞ to +∞) of the *u* variables for smoother gradient ascent, while the constraints on the real parameters are always respected. We propagate the gradient to those variables with the chain rule:

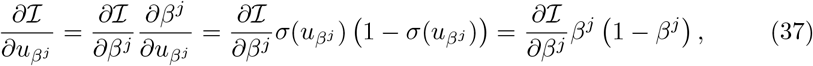

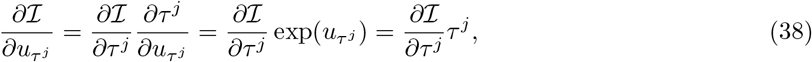

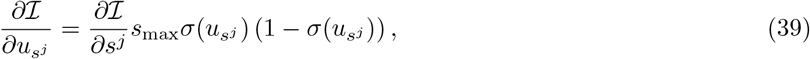

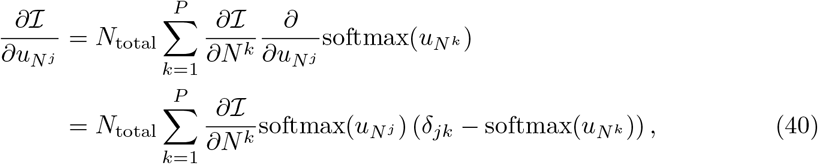

where *δ*_*jk*_ is the Kronecker delta.

For numerical stability, we make sure *β* ∈ [0, 0.99], *τ, N > ϵ* and *s* ∈ [0, 0.99*s*_max_], where *ϵ* = 10^−12^. Note the use of *s*_max_ is not for numerical stability per se. Rather, this value is given by the maximum value of reachable resources *x*_*n*_ for a sequence of length *n*. In other words,

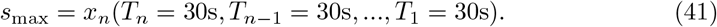

This insures the optimizer never lets a population reach a state where, no matter the sequence of intervals, there is no event-induced response because *s > x*_*n*_ and, thus, the derivative w.r.t. *s* is null.

Retrieving the optimal set of parameters *β, τ, s*, and *N* for a given sequence length *n*, access to recalls through WM with forgetting rate *τ*_f_ and number of populations *P* is done in a few stages.

We first initialize *P* homogeneous populations of adaptive neurons, where each population has a value of *β* ∈ [0, 0.5] and *τ* ∈ [10, 20] sec chosen randomly in their respective intervals. When RT is available, *s* is also uniformly distributed between 0 and 0.5. Each population starts with the same number of neurons and the total number of neurons *N*_total_ is initialized s.t. the random network reaches an MI content of 0.8*H*(**T**), given by Equation (16). All neurons have the same gain Λ = 25 and baseline firing rate Λ_0_ = 5.

The total number of neurons *N*_total_ is held constant during an ascent step. The use of the softmax function lets the optimizer find the right proportions of *N*_total_ shared between the different populations of neurons. However, we wish to stay near an MI value 0.8*H*(**T**) during optimization to make sure the gradient goes to 0 due to reaching the optimum and not because the MI is close to *H*(**T**). To do so, we simply exploit the fact that MI is monotone increasing as a function of *N*_total_ (see Figure 3a). As such, after a gradient step, we update *N*_total_ ← *N*_total_ exp (*α*), where *α* = 0.8*H*(**T**) − I_current_ is clamped between −0.02 and +0.02 and I_current_ is the value of MI computed during the last gradient ascent step. If I_current_ *>* 0.8*H*(**T**), then *N*_total_ is reduced, in turn reducing the value of MI towards 0.8*H*(**T**). Conversely, if I_current_ *<* 0.8*H*(**T**), *N*_total_ is increased. Holding the network near 0.8*H*(**T**) while the softmax changes the proportion of *N*_total_ each population has means the optimizer is not maximizing MI in absolute terms, but achieving the target of 0.8*H*(**T**) with the fewest total number of cells possible. Indeed, if an ascent step makes the MI larger, then *N*_total_ will be reduced on the next step, making the network encode the same amount of information with fewer neurons.

We do a total of 5, 000 gradient ascent steps, each one with a total of *S* = 100, 000 samples to estimate the MI and its gradient. Because the Adam optimizer has an adaptive step size, the small gradient noise near the optima can still generate oscillations of the parameters at the end of the optimization. Therefore, we take the average of the last 500 steps as the optimal values of the parameters. This assumes the optimum is reached within 4, 500 steps and that the optimizer is driven solely by the small gradient noise for the last 500 steps, which appears to be the case in our simulations. An animation of the optimization process is available as Supplementary Animation 1.

## Acknowledgements

This research was enabled in part by support provided by Calcul Québec (www.calculquebec.ca) and the Digital Research Alliance of Canada (alliancecan.ca). This work was supported by NSERC grant RGPIN/06204-2014 to AL and by FRQ grant B2X/328560 to RLM.

## Author contributions

RLM did the theoretical work, mathematical derivations and numerical simulations with input from and verification by AL. LM provided biological background. RLM and AL wrote the manuscript. LM reviewed and edited the manuscript.

## Competing interests

The authors declare no competing interests.

## Data availability

The simulation data necessary to reproduce the figures in this article is available at https://doi.org/10.5281/zenodo.22676828.

## Code availability

The necessary code to reproduce the presented results is available at https://github.com/raphlaf/EfficientAdaptation.

## Notes

### Competing Interest Statement

The authors have declared no competing interest.

