## Supplementary information for "Clustered heterogeneity, spiking history and efficient silencing maximize mutual information in adaptive time encoding"

### 5 Supplementary information

#### Supplementary figures

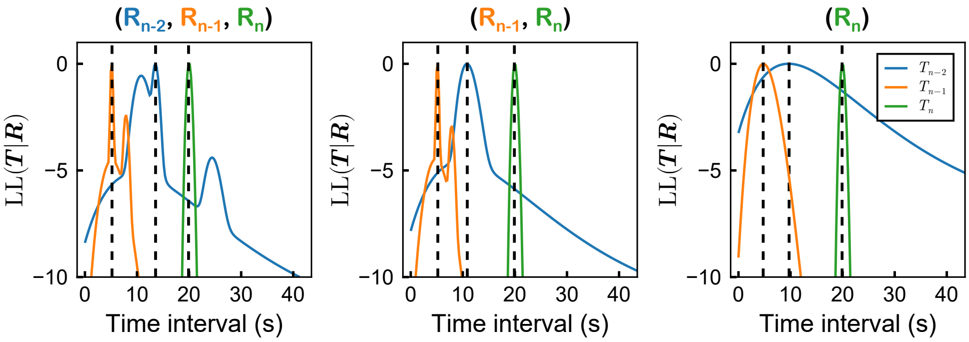


**Supp. Fig. 1**: **Estimation process of time sequences can be irregular.** Loglikelihood (LL) of time interval values in a sequence $\mathbf{T}$ given an observed set of responses and recalls $\mathbf{R}$ from 5 different populations. The situation is identical to what is shown in Figure 2c, with the exception that the sampled responses and recalls are different. Doing maximum likelihood estimation (MLE) on these samples requires taking the maximum value of LL as an estimate for time interval values $(T_{n-2}, T_{n-1}, T_n)$, represented by the dashed lines. Due to the nonlinearity caused by the response threshold and the mixture distribution of the recalled responses, the LL can have an erratic shape with different maxima that are difficult to distinguish from each other, making the estimation less accurate. In the example here, the real sequence of intervals is $(T_{n-2} = 10,\ T_{n-1} = 5,\ T_n = 20)$ s. When adding the recall of $\mathbf{R}_{n-2}$, the LL as a function of $T_{n-2}$ gains two new local maxima, one of which is the global maximum (i.e. the solution) which is further away from the real value of $T_{n-2}$ than the solution given by only $(\mathbf{R}_{n-1}, \mathbf{R}_n)$. This is an example of a possibly biased solution.

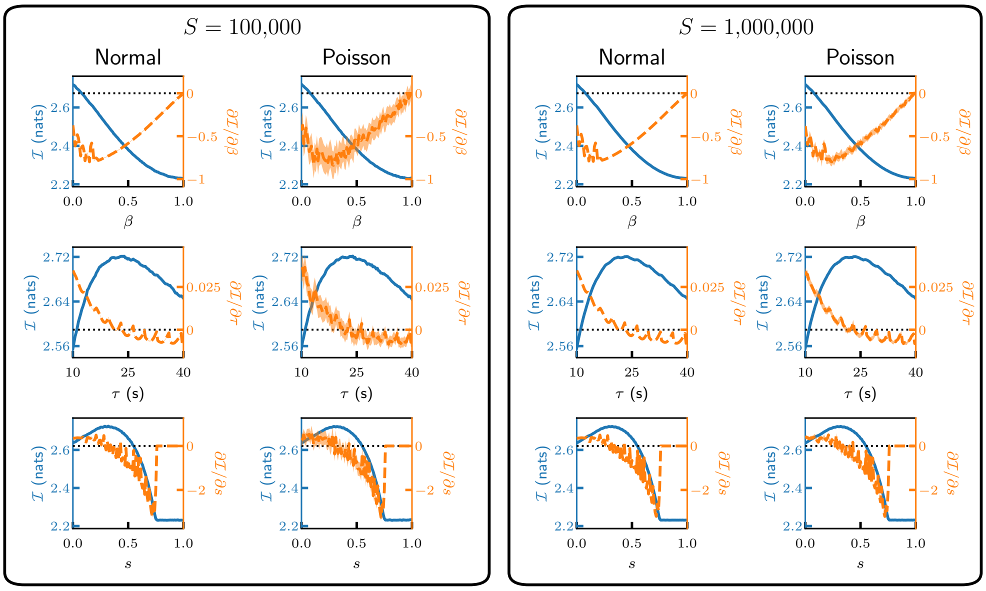


**Supp. Fig. 2**: **Parameter sweep of mutual information and its gradient for Poisson responses and their Normal approximation.** The parameters $\beta$, $\tau$ and $s$ (from top to bottom) of a population among 3 in a network of adaptive neurons are changed while the mutual information (MI, blue solid line) and its derivative (orange dashed line) w.r.t. the changing parameter are estimated using Equations (26), (30) for Poisson populations and (31) for Normal populations. The standard deviation from 10 repetitions of the process is shown in the shaded area and is visible only for the Poisson estimator, though going from $S = 100,000$ samples to $S = 1,000,000$ shrinks the variability significantly. The shape of both estimators agree, which is a sign that the Normal approximation is correct, at least in this regime where $\Lambda_0^j = 5$ for all neurons.

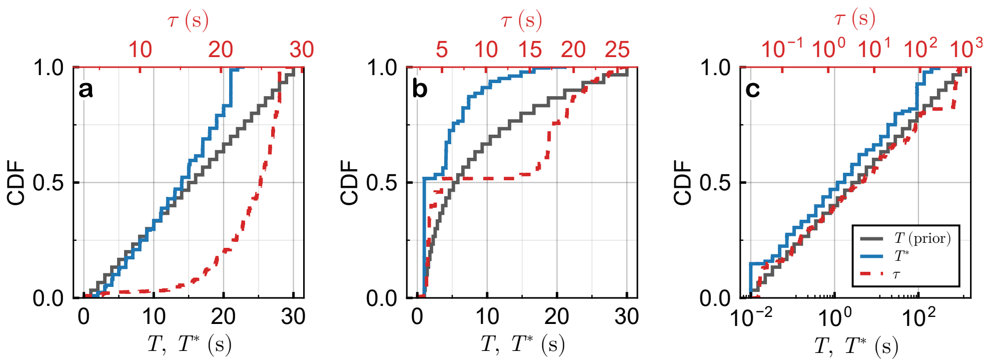


**Supp. Fig. 3**: **A higher objective necessitates higher thresholds.** Same simulation as in Figure 8 with a higher objective MI of $0.99H(\mathbf{T})$ rather than $0.8H(\mathbf{T})$. **(a)** Uniform prior spanning interval values between 1 s and 30 s. **(b)** Powerlaw prior ($P(T) \propto T^{-1}$)) spanning interval values between 1 s and 30 s. **(c)** Powerlaw prior spanning interval values between 0.01 s and 1000 s. The resulting distribution of thresholds retain the same general shape, but reaches higher values of $T^*$ than in Figure 8. Legend is shared across panels.

#### Supplementary methods

##### Sampling from a mixture distribution

While sampling from a mixture distribution numerically is quite simple via the composition method, it is worth laying out explicitly. It especially helps understand how gradients propagate through the sampler itself in the gradient estimator given by Equation (31). First, because all responses and recalls are conditionally independent, we can sample their values independently given the same sample $\mathbf{T}_i$. As such, we look at the sampling of the probability mass function (PMF) or probability density function (PDF) given by Equation (8) for Poisson or Normal responses/recalls, respectively, and repeat the process for all responses and recalls as necessary. For convenience, we rewrite Equation (8) here:

$$P\left(\left\{R^j_{n-m}\right\}_{j=1}^{P} \middle| \{T_k\}_{k=1}^{n}\right) = w_{n-m} \prod_{j=1}^{P} \begin{cases} \frac{1}{\Lambda^j_{\max}+1} & \text{if} R^j_{n-m} \leq \Lambda^j_{\max} \\ 0 & \text{otherwise.} \end{cases}$$
$$+ (1 - w_{n-m}) \prod_{j=1}^{P} \frac{\left(\lambda^j_{n-m}\right)^{R^j_{n-m}} e^{-\lambda^j_{n-m}}}{R^j_{n-m}!}.$$

With only two components in the mixture, we first need to sample a Bernoulli random variable to "decide" which component to draw from, i.e. draw $k_{i,n-m} \in \{0, 1\}$ from

$$P(k_{i,n-m}) = w_{n-m}^{k_{i,n-m}} (1 - w_{n-m})^{1-k_{i,n-m}}. \tag{S-1}$$

If $k_{i,n-m} = 1$, then we draw $R^j_{i,n-m} \in [0, \Lambda^j_{\max}]$ for all $j = 1, ..., P$ from the uniform distribution:

$$P(R^j_{i,n-m}|k_{i,n-m}=1) = \frac{1}{\Lambda^j_{\max}+1}, \tag{S-2}$$

where the normalization constant needs to be changed to $\Lambda^j_{\max}$ in the case of Normal recalls. In the case where $k_{i,n-m} = 0$, we instead draw $R^j_{i,n-m} \in [0, \infty)$ for all $j = 1, ..., P$ from either the Poisson distribution

$$P(R^j_{i,n-m}|k_{i,n-m}=0) = \frac{e^{-\lambda^j_{n-m}}\left(\lambda^j_{n-m}\right)^{R^j_{i,n-m}}}{\left(R^j_{i,n-m}\right)!}, \tag{S-3}$$

or the Normal distribution ($R^j_{i,n-m} \in (-\infty, \infty)$)

$$P(R^j_{i,n-m}|k_{i,n-m}=0) = \frac{\exp\left[-\frac{\left(R^j_{i,n-m}-\lambda^j_{n-m}\right)^2}{2\lambda^j_{n-m}}\right]}{\sqrt{2\pi\lambda^j_{n-m}}}. \tag{S-4}$$

Note how the sampling of the Bernoulli random variable $k_{i,n-m}$ is shared across all populations: all of the populations' recalls are sampled from either their respective uniform or Poisson/Normal distributions.

For the reparameterization trick in the case of Normal responses and recalls, we can define a sampled value as a function of 3 random variables independent of the model parameters. Let $k_{i,n-m} \in \{0, 1\}$ be drawn from the Bernoulli distribution (S-1), $u^j_{i,n-m} \in [0, 1]$ be drawn from a uniform distribution between 0 and 1 and $\epsilon^j_{i,n-m}$ be drawn from a Normal distribution with mean 0 and variance 1. We can then define

$$\begin{aligned} R^j_{i,n-m} = \ & k_{i,n-m} u^j_{i,n-m} \Lambda^j_{\max} \\ & + (1-k_{i,n-m}) \left(\lambda^j_{i,n-m} + \sqrt{\lambda^j_{i,n-m}} \epsilon^j_{i,n-m}\right), \end{aligned} \tag{S-5}$$

where $\lambda^j_{i,n-m}$ is evaluated at the sampled sequence $\mathbf{T}_i$. The derivative of the response w.r.t. a parameter $\theta^j$ is then simply given by

$$\partial_{\theta^j} R^j_{i,n-m} = k_{i,n-m} w^j_{i,n-m} \partial_{\theta^j} \Lambda^j_{\max} + (1 - k_{i,n-m}) \left(1 + \frac{\epsilon^j_{i,n-m}}{2\sqrt{\lambda^j_{i,n-m}}}\right) \partial_{\theta^j} \lambda^j_{i,n-m}. \quad \text{(S-6)}$$

#### Gradient estimation for Poisson populations

Here we derive Equation (30) from the general formulation of MI given in Equation (15) which we rewrite here for convenience:

$$\mathcal{I}(\mathbf{R};\mathbf{T}) = \sum_{(\mathbf{R},\mathbf{T})} P(\mathbf{R}|\mathbf{T})P(\mathbf{T}) \log \left[ \frac{P(\mathbf{R}|\mathbf{T})}{\sum_{\mathbf{T}'} P(\mathbf{R}|\mathbf{T}')P(\mathbf{T}')} \right].$$

First, note that

$$\partial_{\theta^j} \log P(\mathbf{R}|\mathbf{T}) = \frac{\partial_{\theta^j} P(\mathbf{R}|\mathbf{T})}{P(\mathbf{R}|\mathbf{T})} \quad \text{(S-7)}$$

or, equivalently,

$$\partial_{\theta^j} P(\mathbf{R}|\mathbf{T}) = P(\mathbf{R}|\mathbf{T}) \partial_{\theta^j} \log P(\mathbf{R}|\mathbf{T}). \quad \text{(S-8)}$$

We then rewrite Equation (15) as

$$\mathcal{I}(\mathbf{R};\mathbf{T}) = \sum_{(\mathbf{R},\mathbf{T})} \left[ P(\mathbf{R}|\mathbf{T})P(\mathbf{T}) \log P(\mathbf{R}|\mathbf{T}) - P(\mathbf{R}|\mathbf{T})P(\mathbf{T}) \log \left( \sum_{\mathbf{T}'} P(\mathbf{R}|\mathbf{T}')P(\mathbf{T}') \right) \right]. \quad \text{(S-9)}$$

Assigning $I_1$ to the first term and $I_2$ to the second one, we compute the derivatives of both terms separately. For $I_1$, we use Equation (S-8) to get

$$
\begin{aligned}
\partial_{\theta^j} I_1 &= \sum_{(\mathbf{R},\mathbf{T})} P(\mathbf{T}) \partial_{\theta^j} \left[ P(\mathbf{R}|\mathbf{T}) \log P(\mathbf{R}|\mathbf{T}) \right] \\
&= \sum_{(\mathbf{R},\mathbf{T})} P(\mathbf{T}) \left[ \log P(\mathbf{R}|\mathbf{T}) \partial_{\theta^j} P(\mathbf{R}|\mathbf{T}) + P(\mathbf{R}|\mathbf{T}) \partial_{\theta^j} \log P(\mathbf{R}|\mathbf{T}) \right] \\
&= \sum_{(\mathbf{R},\mathbf{T})} P(\mathbf{T}) \left[ \log P(\mathbf{R}|\mathbf{T}) P(\mathbf{R}|\mathbf{T}) \partial_{\theta^j} \log P(\mathbf{R}|\mathbf{T}) + P(\mathbf{R}|\mathbf{T}) \partial_{\theta^j} \log P(\mathbf{R}|\mathbf{T}) \right] \\
&= \sum_{(\mathbf{R},\mathbf{T})} P(\mathbf{T}) \left[ \log P(\mathbf{R}|\mathbf{T}) + 1 \right] P(\mathbf{R}|\mathbf{T}) \partial_{\theta^j} \log P(\mathbf{R}|\mathbf{T}). \qquad \text{(S-10)}
\end{aligned}
$$

For the second term, we have

$$
\begin{aligned}
\partial_{\theta^j} I_2 &= \sum_{\mathbf{R}} \sum_{\mathbf{T}} P(\mathbf{T}) \partial_{\theta^j} \left[ P(\mathbf{R}|\mathbf{T}) \log \left( \sum_{\mathbf{T}'} P(\mathbf{R}|\mathbf{T}') P(\mathbf{T}') \right) \right] \\
&= \sum_{\mathbf{R}} \sum_{\mathbf{T}} P(\mathbf{T}) \partial_{\theta^j} P(\mathbf{R}|\mathbf{T}) \log \left( \sum_{\mathbf{T}'} P(\mathbf{R}|\mathbf{T}') P(\mathbf{T}') \right) \\
&\quad + \sum_{\mathbf{R}} \sum_{\mathbf{T}} P(\mathbf{T}) P(\mathbf{R}|\mathbf{T}) \frac{\partial_{\theta^j} \left[ \sum\limits_{\mathbf{T}'} P(\mathbf{R}|\mathbf{T}') P(\mathbf{T}') \right]}{\sum\limits_{\mathbf{T}''} P(\mathbf{R}|\mathbf{T}'') P(\mathbf{T}'')} \\
&= \sum_{\mathbf{R}} \sum_{\mathbf{T}} P(\mathbf{T}) P(\mathbf{R}|\mathbf{T}) \partial_{\theta^j} \log P(\mathbf{R}|\mathbf{T}) \log \left( \sum_{\mathbf{T}'} P(\mathbf{R}|\mathbf{T}') P(\mathbf{T}') \right) \\
&\quad + \sum_{\mathbf{R}} \frac{\sum\limits_{\mathbf{T}'} P(\mathbf{T}') \partial_{\theta^j} P(\mathbf{R}|\mathbf{T}')}{\sum\limits_{\mathbf{T}''} P(\mathbf{R}|\mathbf{T}'') P(\mathbf{T}'')} \sum_{\mathbf{T}} P(\mathbf{R}|\mathbf{T}) P(\mathbf{T}) \\
&= \sum_{\mathbf{R}} \sum_{\mathbf{T}} P(\mathbf{T}) P(\mathbf{R}|\mathbf{T}) \partial_{\theta^j} \log P(\mathbf{R}|\mathbf{T}) \log \left( \sum_{\mathbf{T}'} P(\mathbf{R}|\mathbf{T}') P(\mathbf{T}') \right) \\
&\quad + \sum_{\mathbf{R}} \cancelto{1}{\frac{\sum\limits_{\mathbf{T}} P(\mathbf{R}|\mathbf{T}) P(\mathbf{T})}{\sum\limits_{\mathbf{T}''} P(\mathbf{R}|\mathbf{T}'') P(\mathbf{T}'')}} \sum_{\mathbf{T}'} P(\mathbf{T}') \partial_{\theta^j} P(\mathbf{R}|\mathbf{T}') \\
&= \sum_{\mathbf{R}} \sum_{\mathbf{T}} P(\mathbf{T}) P(\mathbf{R}|\mathbf{T}) \partial_{\theta^j} \log P(\mathbf{R}|\mathbf{T}) \log \left( \sum_{\mathbf{T}'} P(\mathbf{R}|\mathbf{T}') P(\mathbf{T}') \right) \\
&\quad + \sum_{\mathbf{R}} \sum_{\mathbf{T}'} P(\mathbf{T}') P(\mathbf{R}|\mathbf{T}') \partial_{\theta^j} \log P(\mathbf{R}|\mathbf{T}') \\
&= \sum_{(\mathbf{R},\mathbf{T})} \left[ \log \left( \sum_{\mathbf{T}'} P(\mathbf{R}|\mathbf{T}') P(\mathbf{T}') \right) + 1 \right] P(\mathbf{T}) P(\mathbf{R}|\mathbf{T}) \partial_{\theta^j} \log P(\mathbf{R}|\mathbf{T}). \qquad \text{(S-11)}
\end{aligned}
$$

We then combine Equations (S-10) and (S-11) to get

$$
\partial_{\theta^j} \mathcal{I} = \sum_{(\mathbf{R},\mathbf{T})} P(\mathbf{R}|\mathbf{T}) P(\mathbf{T}) \left[ \log \left( \frac{P(\mathbf{R}|\mathbf{T})}{\sum_{\mathbf{T}'} P(\mathbf{R}|\mathbf{T}') P(\mathbf{T}')} \right) \right] \partial_{\theta^j} \log P(\mathbf{R}|\mathbf{T}). \qquad \text{(S-12)}
$$

We can also note that, with Equation (S-7), we have

$$
\begin{aligned}
\sum_{(\mathbf{R},\mathbf{T})} P(\mathbf{R}|\mathbf{T})P(\mathbf{T})\partial_{\theta^j}\log P(\mathbf{R}|\mathbf{T}) &= \sum_{(\mathbf{R},\mathbf{T})} \frac{P(\mathbf{R}|\mathbf{T})}{P(\mathbf{R}|\mathbf{T})} P(\mathbf{T})\partial_{\theta^j} P(\mathbf{R}|\mathbf{T}) \\
&= \partial_{\theta^j}\left[\sum_{(\mathbf{R},\mathbf{T})} P(\mathbf{R}|\mathbf{T})P(\mathbf{T})\right] \\
&= \partial_{\theta^j}[1] = 0. \qquad \text{(S-13)}
\end{aligned}
$$

As such, we can add or subtract any constant (independent of $\mathbf{R}$) inside the square brackets of Equation (S-12) and still retrieve the same value in the end. This allows the use of a control variate that reduces the variance of a Monte Carlo estimator of Equation (S-12) without biasing the final value. This is what is done in Equation (30) in the main article where the control variate is the current estimate of the MI.

#### Gradient estimation for Normal populations

When the homogeneous populations of neurons are large enough and have a significant baseline, the minimum firing parameter being $N^j\Lambda_0^j$ means that the average of the distribution of the sum of spikes of said population is always far from 0. As such, we can approximate this Poisson distribution as a Normal distribution. We can then redefine the sample $R^j_{i,n-m}$ as in Equation (S-5). As such, if we rewrite the MI as an expectation value as

$$
\mathcal{I}(\mathbf{R};\mathbf{T}) = \underset{(\mathbf{R},\mathbf{T})}{\mathrm{E}}\left[\log\left(\frac{P(\mathbf{R}|\mathbf{T})}{\underset{\mathbf{T}'}{\mathrm{E}}\left[P(\mathbf{R}|\mathbf{T}')\right]}\right)\right] \qquad \text{(S-14)}
$$

and note that taking the expectation over $\mathbf{R}$ is now equivalent to taking the expectation over $\boldsymbol{\omega}$ — i.e. the set of all random variables independent of optimized parameters $\left\{\epsilon^j_{n-m}, u^j_{n-m}, k_{n-m}\right\}_{m=0,j=1}^{m=n-1,j=P}$ — we get

$$\mathcal{I}(\mathbf{R};\mathbf{T}) = \mathop{\mathrm{E}}_{(\boldsymbol{\omega},\mathbf{T})}\left[\log\left(\frac{P(\mathbf{R}(\boldsymbol{\omega})|\mathbf{T})}{\mathop{\mathrm{E}}_{\mathbf{T}'}\left[P(\mathbf{R}(\boldsymbol{\omega})|\mathbf{T}')\right]}\right)\right]. \tag{S-15}$$

This expression means that we can now take the derivative of $\mathcal{I}$ quite simply as

$$\partial_{\theta^j}\mathcal{I}(\mathbf{R};\mathbf{T}) = \mathop{\mathrm{E}}_{(\boldsymbol{\omega},\mathbf{T})}\left[\partial_{\theta^j}\log\left(\frac{P(\mathbf{R}(\boldsymbol{\omega})|\mathbf{T})}{\mathop{\mathrm{E}}_{\mathbf{T}'}\left[P(\mathbf{R}(\boldsymbol{\omega})|\mathbf{T}')\right]}\right)\right], \tag{S-16}$$

as the sampling, and hence the expectation itself, no longer depends on the parameters $\theta^j$. All that remains is to explicitly take the derivatives of what is inside the square brackets in Equation (S-16)

$$\begin{aligned}
\partial_{\theta^j}\mathcal{I}(\mathbf{R};\mathbf{T}) &= \mathop{\mathrm{E}}_{(\boldsymbol{\omega},\mathbf{T})}\left[\partial_{\theta^j}\log P(\mathbf{R}(\boldsymbol{\omega})|\mathbf{T}) - \frac{\mathop{\mathrm{E}}_{\mathbf{T}'}\left[\partial_{\theta^j}P(\mathbf{R}(\boldsymbol{\omega})|\mathbf{T}')\right]}{\mathop{\mathrm{E}}_{\mathbf{T}''}\left[P(\mathbf{R}(\boldsymbol{\omega})|\mathbf{T}'')\right]}\right] \\
&= \mathop{\mathrm{E}}_{(\boldsymbol{\omega},\mathbf{T})}\Bigg[\partial_{\theta^j}\log P(\mathbf{R}(\boldsymbol{\omega})|\mathbf{T}) \\
&\qquad - \mathop{\mathrm{E}}_{\mathbf{T}'}\left[\frac{P(\mathbf{R}(\boldsymbol{\omega})|\mathbf{T}')}{P(\mathbf{R}(\boldsymbol{\omega}))}\partial_{\theta^j}\log P(\mathbf{R}(\boldsymbol{\omega})|\mathbf{T}')\right]\Bigg] \\
&= \mathop{\mathrm{E}}_{(\boldsymbol{\omega},\mathbf{T})}\Bigg[\partial_{\theta^j}\log P(\mathbf{R}(\boldsymbol{\omega})|\mathbf{T}) \\
&\qquad - \mathop{\mathrm{E}}_{\mathbf{T}'}\left[\frac{P(\mathbf{T}'|\mathbf{R}(\boldsymbol{\omega}))}{P(\mathbf{T}')}\partial_{\theta^j}\log P(\mathbf{R}(\boldsymbol{\omega})|\mathbf{T}')\right]\Bigg],
\end{aligned} \tag{S-17}$$

where we used Equation (S-8) in the second step and Bayes' theorem in the third step. Equation (S-17) is what is being computed with Monte Carlo simulations in Equation (31).

#### Derivatives of log-conditionals

Here we derive the general equation for $\partial_{\theta^j}\log P(\mathbf{R}_i|\mathbf{T}')$ for any pair of $\mathbf{R}_i \sim P(\mathbf{R}|\mathbf{T}_i)$ and sequence $\mathbf{T}'$. We first define $U^j_{n-m}(R^j_{i,n-m})$ as the uniform probability distribution of the recall of the $n-m$-th interval of population $j$ and $Q^j_{n-m}(R^j_{i,n-m}, T'_1, ...T'_{n-m})$ as either the Poisson or Normal probability distribution of the current response ($m = 0$) or of the recall of the $n-m$-th interval of population $j$ evaluated at sequence $\mathbf{T}'$. We can then rewrite Equation (8) for the full set of responses and recalls as

$$P(\mathbf{R}_i|\mathbf{T}') = \prod_{m=0}^{n-1} \left[ w'_{n-m} \prod_{j=1}^{P} U^j_{n-m} + (1 - w'_{n-m}) \prod_{j=1}^{P} Q^j_{n-m} \right], \tag{S-18}$$

where $w'_{n-m} = (T'_{n-m+1} + \cdots + T'_n)/\tau_\mathrm{f}$.

We take the derivative of $\log P(\mathbf{R}_i|\mathbf{T}')$ w.r.t. any parameter $\theta^j = \beta^j, \tau^j, s^j, N^j, \Lambda^j, \Lambda^j_0$ of population $j$ as (dropping the explicit dependencies on the $\mathbf{R}$ and $\mathbf{T}$ for a lighter notation)

$$\begin{aligned}
\partial_{\theta^j} \log P(\mathbf{R}_i|\mathbf{T}') &= \sum_{m=0}^{n-1} \partial_{\theta^j} \log \left[ w'_{n-m} \prod_{j'=1}^{P} U^{j'}_{n-m} + (1 - w'_{n-m}) \prod_{j'=1}^{P} Q^{j'}_{n-m} \right] \\
&= \sum_{m=0}^{n-1} \frac{\left[ w'_{n-m} \partial_{\theta^j} \left( \prod_{j'=1}^{P} U^{j'}_{n-m} \right) + (1 - w'_{n-m}) \partial_{\theta^j} \left( \prod_{j'=1}^{P} Q^{j'}_{n-m} \right) \right]}{\left[ w'_{n-m} \prod_{j'=1}^{P} U^{j'}_{n-m} + (1 - w'_{n-m}) \prod_{j'=1}^{P} Q^{j'}_{n-m} \right]} \\
&= \sum_{m=0}^{n-1} \frac{w'_{n-m} \prod_{j'=1, j'\neq j}^{P} U^{j'}_{n-m} \partial_{\theta^j} U^j_{n-m}}{\left[ w'_{n-m} \prod_{j'=1}^{P} U^{j'}_{n-m} + (1 - w'_{n-m}) \prod_{j'=1}^{P} Q^{j'}_{n-m} \right]} \\
&\quad + \sum_{m=0}^{n-1} \frac{(1 - w'_{n-m}) \prod_{j'=1, j'\neq j}^{P} Q^{j'}_{n-m} \partial_{\theta^j} Q^j_{n-m}}{\left[ w'_{n-m} \prod_{j'=1}^{P} U^{j'}_{n-m} + (1 - w'_{n-m}) \prod_{j'=1}^{P} Q^{j'}_{n-m} \right]} \\
&= \sum_{m=0}^{n-1} \frac{w'_{n-m} \prod_{j'=1}^{P} U^{j'}_{n-m} \partial_{\theta^j} \log U^j_{n-m}}{\left[ w'_{n-m} \prod_{j'=1}^{P} U^{j'}_{n-m} + (1 - w'_{n-m}) \prod_{j'=1}^{P} Q^{j'}_{n-m} \right]} \\
&\quad + \sum_{m=0}^{n-1} \frac{(1 - w'_{n-m}) \prod_{j'=1}^{P} Q^{j'}_{n-m} \partial_{\theta^j} \log Q^j_{n-m}}{\left[ w'_{n-m} \prod_{j'=1}^{P} U^{j'}_{n-m} + (1 - w'_{n-m}) \prod_{j'=1}^{P} Q^{j'}_{n-m} \right]} \\
&= \sum_{m=0}^{n-1} \left[ (1 - \gamma_{n-m}) \partial_{\theta^j} \log U^j_{n-m} + \gamma_{n-m} \partial_{\theta^j} \log Q^j_{n-m} \right],
\end{aligned} \tag{S-19}$$

where

$$\gamma_{n-m} = \frac{(1-w'_{n-m})\prod_{j'=1}^{P} Q_{n-m}^{j'}}{w'_{n-m}\prod_{j'=1}^{P} U_{n-m}^{j'} + (1-w'_{n-m})\prod_{j'=1}^{P} Q_{n-m}^{j'}} \tag{S-20}$$

is a value between 0 and 1 indicating how much the Poisson/Normal part of the mixture contributes to the value of the derivative.

We can now compute the derivatives of the uniform and Poisson/Normal parts of each population separately. First, in the case of **Poisson** responses and recalls, we have

$$\partial_{\theta^j} \log U_{n-m}^j = \begin{cases} -\frac{\partial_{\theta^j} \Lambda_{\max}^j}{\Lambda_{\max}^j + 1} & \text{if} \quad 0 \leq R_{i,n-m}^j \leq \Lambda_{\max}^j \\ 0 & \text{otherwise,} \end{cases} \tag{S-21}$$

and

$$\partial_{\theta^j} \log Q_{n-m}^j = \left( \frac{R_{i,n-m}^j}{\lambda_{n-m}^{'j}} - 1 \right) \partial_{\theta^j} \lambda_{n-m}^{'j}, \tag{S-22}$$

where $\lambda_{n-m}^{'j} = \lambda_{n-m}^j(T'_1, ..., T'_{n-m})$ is the firing parameter (2) at any sequence $\mathbf{T}'$.

The **Normal** responses and recalls are similar, with the exception that we take into account the derivative given by Equation (S-6). We thus have

$$\partial_{\theta^j} \log U_{n-m}^j = \begin{cases} -\frac{\partial_{\theta^j} \Lambda_{\max}^j}{\Lambda_{\max}^j} & \text{if} \quad 0 \leq R_{i,n-m}^j \leq \Lambda_{\max}^j \\ 0 & \text{otherwise,} \end{cases} \tag{S-23}$$

and

$$\begin{aligned} \partial_{\theta^j} \log Q_{n-m}^j = -\frac{1}{2(\lambda_{n-m}^{'j})^2} \Bigg[ &\left( \lambda_{n-m}^{'j} + \left(\lambda_{n-m}^{'j}\right)^2 - \left(R_{i,n-m}^j\right)^2 \right) \partial_{\theta^j} \lambda_{n-m}^{'j} \\ &+ 2\lambda_{n-m}^{'j} \left( R_{i,n-m}^j - \lambda_{n-m}^{'j} \right) \partial_{\theta^j} R_{i,n-m}^j \Bigg]. \end{aligned} \tag{S-24}$$

Note that the firing parameter $\lambda'^{j}_{n-m}$ and its derivative in Equation (S-24) is evaluated at a sequence $\mathbf{T}'$, not necessarily the sampled sequence. However, the firing parameter $\lambda^{j}_{n-m}$ and its derivative coming from Equation (S-6) in the second term of Equation (S-24) is evaluated at the sampled sequence $\mathbf{T}_i$.

The derivatives of $\lambda$ and $\Lambda_{\max}$ are shared by both the Poisson and the Normal populations. For $\Lambda_{\max}$, we have (we drop the $j$ indices for a lighter notation)

$$\partial_N \Lambda_{\max} = \Lambda + \Lambda_0 + 5\sqrt{\frac{\Lambda + \Lambda_0}{N}}, \tag{S-25}$$

$$\partial_\Lambda \Lambda_{\max} = \partial_{\Lambda_0} \Lambda_{\max} = N + 5\sqrt{\frac{N}{\Lambda + \Lambda_0}}, \tag{S-26}$$

with the derivatives w.r.t. all other parameters being 0.

For the firing parameter, we have

$$\partial_\beta \lambda_{n-m} = N\Lambda \partial_\beta x_{n-m}, \tag{S-27}$$

$$\partial_\tau \lambda_{n-m} = N\Lambda \partial_\tau x_{n-m}, \tag{S-28}$$

$$\partial_s \lambda_{n-m} = -N\Lambda, \tag{S-29}$$

$$\partial_\Lambda \lambda_{n-m} = N[x_{n-m} - s]_+, \tag{S-30}$$

all dependent on $x_{n-m} > s$. Otherwise these values are 0. Then we have

$$\partial_N \lambda_{n-m} = \Lambda[x_{n-m} - s]_+ + \Lambda_0, \tag{S-31}$$

$$\partial_{\Lambda_0} \lambda_{n-m} = N. \tag{S-32}$$

Finally, we compute the derivatives of the resource variable $x$ w.r.t. $\beta$ and $\tau$ recursively:

$$\partial_\beta x_n = e^{-T_n/\tau} \left( x_{n-1} + \beta \partial_\beta x_{n-1} \right), \tag{S-33}$$

$$\partial_\tau x_n = e^{-T_n/\tau} \left[ \beta \partial_\tau x_{n-1} - \frac{T_n}{\tau^2} (1 - \beta x_{n-1}) \right], \tag{S-34}$$

with $\partial_\beta x_1 = e^{-T_1/\tau}$ and $\partial_\tau x_1 = -\frac{T_1}{\tau^2} e^{-T_1/\tau}(1-\beta)$.
